# Differential expression analysis of sexual and apomictic *Boechera* uncovers *FAS4* as crucial for gametogenesis

**DOI:** 10.1101/2022.10.05.510110

**Authors:** Laura Binmöller, Christopher Volkert, Christiane Kiefer, Luise Zühl, Magdalena W. Slawinska, Anna Loreth, Berit H. Nauerth, David Ibberson, Rafael Martinez, Reinhard Zipper, Maike Kohnle, Anja Schmidt

## Abstract

During sexual reproduction of higher plants, seed formation is initiated by double fertilization of egg and central cell. In contrast, pseudogamous apomicts form embryos asexually by parthenogenesis of the egg, but initiation of endosperm development still depends on central cell fertilization. It can be envisioned that these differences are determined during gametogenesis and specification of gametophytic cells. To deepen the understanding of the transcriptional basis underlying sexual and apomictic reproduction, we applied tissue type-specific RNA-Seq. We compared expression in reproductive tissues of different *Boechera* accessions at distinct developmental stages. This confirmed previous evidence for an enrichment of RNA helicases at onset of reproductive development. We further identified a small number of members of this gene family as differentially expressed in female reproductive ovule tissues harbouring mature gametophytes from apomictic and sexual accessions. This included homologues of *A. thaliana FASCIATED STEM 4* (*FAS4*) and of *ENHANCED SILENCING PHENOTYPE 3* (*ESP3*), which have previously been identified as potential candidates for gametogenesis and apomixis, respectively. Unlike in *A. thaliana*, for either of them additional homologues or copies of related genes are present in *Boechera*, indicating complex evolutionary histories. As the expression patterns implied potential roles of *FAS4* during gametogenesis, we first studied *A. thaliana* lines carrying mutant alleles. Indeed, we observed defects during male and female gametogenesis and severely reduced transmission efficiencies through both parents. In conclusion, our study identifies *FAS4* as crucial for plant reproduction and suggests the potential for sub-functionalization of additional homologous genes in *Boechera* to shape reproductive development.

## Introduction

In the life cycle of higher plants, the formation of seeds is a key step that mediates the transition from the gametophytic to the next sporophytic generation. During sexual reproduction, double fertilization of egg and central cell with one sperm cell each is required to initiate development of the embryo and its nourishing tissue, the endosperm. Both, formation of sperms as male and of egg and central cells as female gametes progresses in series of defined developmental processes. To initiate pollen formation, a single sporophytic cell, the pollen mother cell, gets selected in the male reproductive flower tissues (anthers). It progresses through meiosis to give rise to a tetrad of microspores. Each microspore subsequently undergoes an asymmetric pollen mitosis I (PMI) to form a generative cell engulfed in a vegetative cell, followed by PMII of the generative cell to give rise to mature pollen (Twell et al., 2006). To mediate the transition from sporophytic to gametophytic fate during female megasporogenesis, first a single sporophytic cell gets selected in the subepidermal tissues of the nucellus domain of the ovule. It specifies into the megaspore mother cell (MMC), which undergoes meiosis to form four megaspores. Of these, typically only the functional megaspore (FMS) survives and serves as founder cell of the gametophytic lineage. Gametogenesis subsequently progresses through three mitotic divisions in a polarized syncytium followed by cellularization to give rise to the mature gametophyte typically consisting of four specialized cell types (Sprunck and Groß-Hardt, 2011). Apart from the female gametes, two synergids, which are important for pollen tube attraction and double fertilization, and antipodal cells are formed.

Understanding the molecular mechanisms governing gametogenesis is not only of great interest in the context of reproduction. Based on its simple organization and short lifespan, the female gametophyte is an attractive model to study organogenesis and cell type specification (Schmid et al., 2015). Even though in the majority of angiosperms gametogenesis is highly conserved, different types of gametophytes are described, allowing evolutionary studies (Friedman and Williams, 2003). Further diversity of reproductive systems is presented, as apart from sexual reproduction, also asexual reproduction through seeds (apomixis) occurs in ∼1% of angiosperms (Hörandl and Hojsgaard, 2012). From a developmental perspective, apomictic development differs from sexual reproduction only in a few steps. Similar to sexual reproduction, the mature gametophyte is formed subsequent to three mitotic divisions and cellularization. But in contrast to sexual reproduction, the first cell of the gametophytic lineage is an uninucleate gametophyte derived from an apomictic initial cell (AIC), which has altered or omitted meiosis to preserve the genetic constitution of the mother plant (Schmidt et al., 2015). Afterwards, the unfertilized egg cell enters embryogenesis (parthenogenesis). Whereas most apomicts require fusion of the central cell nucleus with one sperm to allow endosperm development (pseudogamy), in certain apomicts also endosperm forms autonomously (Barcaccia and Albertini, 2013).

It has long been hypothesized that epigenetic regulatory mechanisms are important in the regulatory control of apomixis (Schmidt, 2020, Grimanelli, 2012, Kumar, 2017). In this line, in sexual species embryo and endosperm development remain repressed prior to fertilization. This involves cell cycle arrest and a repressive transcriptional state of the egg, and for both gametes chromatin modifications and additional epigenetic regulatory mechanisms repressing gene activity based on histone modifications and DNA methylation (Vijverberg et al., 2019). Recently, important new insights into the molecular mechanisms controlling parthenogenesis have been gained (Conner and Ozias-Akins, 2017, Khanday et al., 2019, Underwood et al., 2022). Nevertheless, little is known about the gene regulatory mechanisms controlling pseudogamy. As the egg cell has parthenogenetic fate, but embryogenesis remains repressed prior to central cell fertilization or endosperm development, it is likely that this might be pre-determined during gametogenesis.

To date, knowledge remains limited about the molecular mechanisms governing cell fate acquisition during gametogenesis especially in apomicts. From investigations of sexual reproduction, increasing evidence suggests that hormonal pathways including cytokinin and auxin are involved in establishing polarity during gametogenesis, which in turn is important for cell fate acquisition (Terceros et al., 2020). Migration and localization of the gametophytic nuclei in the syncytial embryo sac is critical in this process (Sprunck and Groß-Hardt, 2011). This was identified based on mutants showing aberrant numbers or positioning of nuclei in developing gametophytes, for example *A. thaliana* mutants of the cell cycle regulator *retinoblastoma-related1* (*rbr1*) (Ebel et al., 2004). Further evidence suggests polar localization of transcripts during gametogenesis (Schmid MW, 2015), which likely involves RNA binding proteins.

In line with this, cell and tissue type-specific transcriptional studies have previously provided important insights into gene regulatory processes underlying reproductive development in *A. thaliana* and accessions of the closely related genus *Boechera* (Schmidt et al., 2014, Zühl et al., 2019, Schmidt et al., 2011, Schmid et al., 2012, Wuest et al., 2010). This indicated the importance of RNA helicases, translational regulation and ribosome biogenesis for the specification of the female germline (Schmidt et al., 2011). Furthermore, based on transcriptional analysis in different species, the RNA helicases *ENHANCED SILENCING PHENOTYPE 3* (*ESP3*) and *MATERNAL EFFECT EMBRYO ARREST 29* (*MEE29*) have been proposed as candidates for apomixis (Barcaccia and Albertini, 2013, Schmidt et al., 2014). Interestingly, in *A. thaliana* both are essential for seed development (Meinke, 2020, Pagnussat et al., 2005). In addition, plants carrying mutant alleles of *MNEME* or *RH17* show features of apomixis (Stein et al., 2021, Schmidt et al., 2011).

In general, RNA helicases form a large and diverse gene family functioning in various aspects of RNA metabolism, including for example splicing, RNA degradation and nonsense mediated decay (Linder and Owttrim, 2009). Furthermore, they are involved in epigenetic regulatory pathways, for example by modification of chromatin structure (Barak et al., 2014). RNA helicases are also frequently associated to ribonucleoprotein complexes (RNP-complexes) important for ribosome assembly and degradation, and for translational control (Van Treeck and Parker, 2018). Amongst the 161 RNA helicases in *A. thaliana* (Xu et al., 2013), 58 DExD/H-box RNA helicases are represented, many of which show homologies to yeast or human genes encoding for ribosomal binding proteins (Mingam et al., 2004, Liu and Imai, 2018). So far, roles for gametogenesis and ribosome biogenesis are described only for a few of them, including *MAGATAMA3* (*MAA3*) and *RH36/SLOW WALKER 3* (*SWA3*) (Huang et al., 2010, Liu et al., 2010, Shimizu et al., 2008). Mutant alleles of *SWA3* lead to a delayed progression through female gametogenesis (Huang et al., 2010, Liu et al., 2010, Shimizu et al., 2008). Defects during mitotic divisions and cell expansion of the syncytial gametophyte have further been demonstrated for lines carrying an RNAi construct for downregulation of the *SWI2/SNF2 CHROMATIN REMODELLING FACTOR 11* (*CHR11*) (Huanca-Mamani et al., 2005). Together, this suggests that several distinct RNA helicases are involved in controlling aspects of gametogenesis.

To gain further insights into the transcriptional regulation underlying germline specification and development in sexual and apomictic *Boechera*, we extended previous tissue type-specific analyses (Zühl et al., 2019). We isolated nucellus tissues harbouring MMC/AICs, or FMS/uninucleate gametophytes at onset of gametogenesis by laser assisted microdissection (LAM), and ovules harbouring mature gametophytes at three days after emasculation by microdissection. Comprehensive RNA-Seq and statistical analysis supported previous evidence for ribosomal pathways, RNA helicases and epigenetic regulatory pathways to play important roles for reproductive development independent of the mode of reproduction. Interestingly, our analysis also identified a few RNA helicases to be differentially regulated in mature ovules from sexual as compared to apomictic accessions. Thereby, homologues of *ESP3, FAS4, CHR34*, and *RH1* were significantly higher expressed in ovules from sexual *Boechera*. Furthermore, we identified that *ESP3* is evolutionary closely related to *MEE29* and to additional so far uncharacterized genes. As *FAS4* is predominantly expressed in synergids in *A. thaliana* (Wuest et al., 2010), it is a promising candidate for gametogenesis. While over-expression of *AtFAS4* leads to stem fasciation (Pogorelko et al., 2008), roles for reproduction have so far not been described. Interestingly, *AtFAS4* is a unique gene, whereas one orthologue and two paralogues are represented in the *Boechera stricta* LTM genome v1.2 (DOE-JGI, http://phytozome.jgi.doe.gov/). To initiate the functional characterization of *FAS4*, we used *A. thaliana* lines carrying mutant alleles of the gene. They showed male and female gametophytic defects and aberrant numbers and positioning of nuclei in female gametophytes. In summary, our study uncovered *AtFAS4* as crucial for gametogenesis and provided new evidence that RNA helicases and their evolutionary history can shape the plasticity of plant reproductive development.

## Results

### Comparative analysis provides comprehensive insights into the transcriptional basis underlying (a)sexual reproduction

Transcriptional analysis of nucellus tissues harbouring MMC or AIC in sexual and apomictic *Boechera* has previously allowed the identification of new candidate genes for specification of the female reproductive lineage in apomicts (Zühl et al., 2019). To gain further insights into these processes and to identify genes potentially involved in gametogenesis, we extended our analysis to nucellus tissues and ovules harbouring FMS/uninucleate gametophytes and mature gametophytes, respectively (Figure 1). A major aim of our study was a closer investigation of the spatial and temporal expression of RNA helicases. Here, seven sexual or apomictic *Boechera* accessions were used (Table 1, Table S1). For nucellus tissues at the two developmental stages, single end (SE) sequencing on the Illumina NextSeq 500 platform generated between 51‘252‘215 and 127‘761‘363 reads of overall good quality (Table S1). Moreover, between 20‘589‘962 and 67‘634‘805 reads were obtained by paired end (PE) sequencing of libraries derived from RNA isolated from ovules (Table S1).

**Figure 1:**
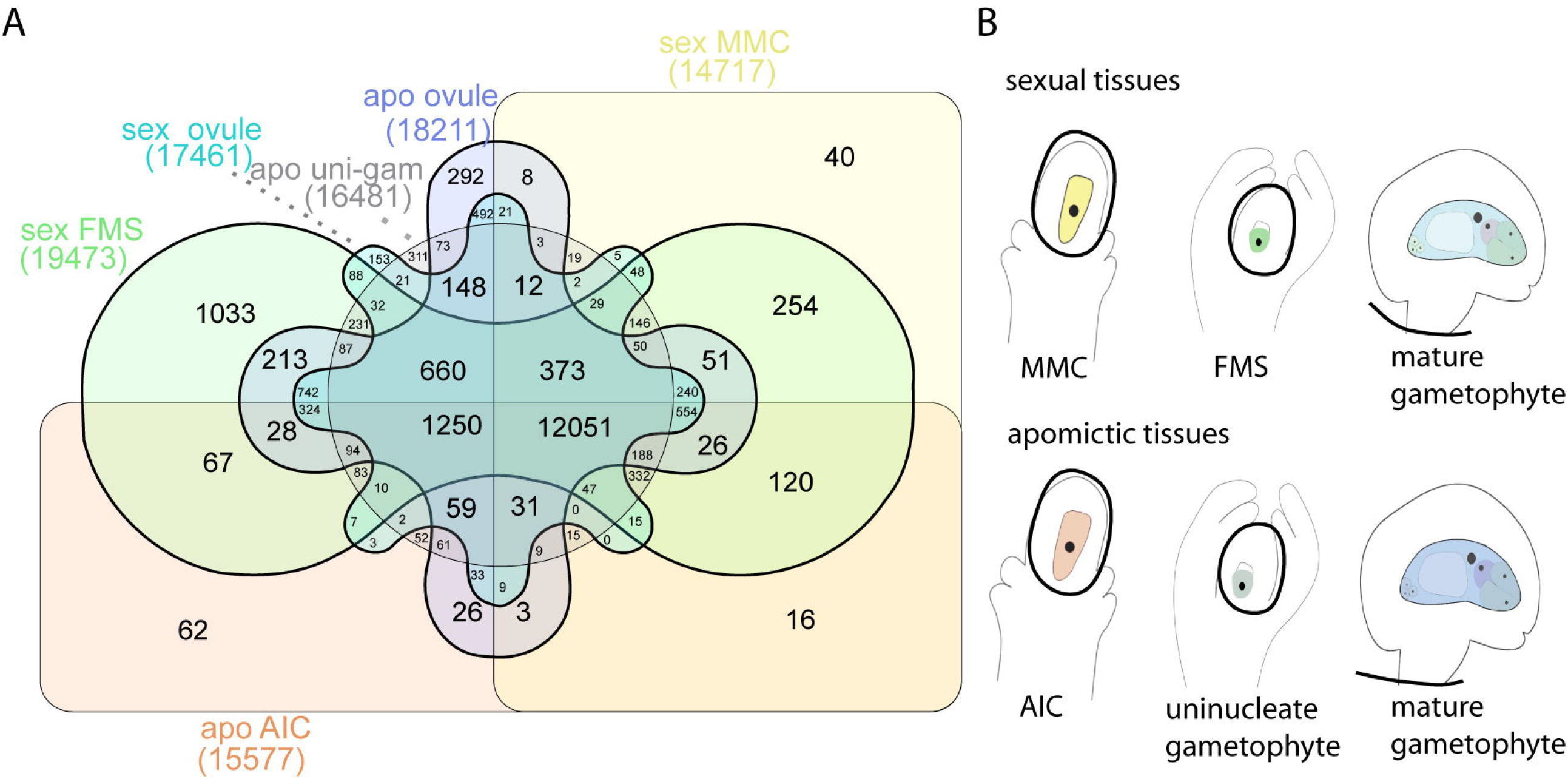
Comparison of gene expression between reproductive modes and developmental stages. (A) Venn diagram showing intersections of gene expression (≥10 TMM normalized read counts) in nucellus tissues harbouring the MMC/AIC, or FMS/uninucleate gametophytes (apo uni-gam, apomictic uninucleate gametophyte), and ovules harbouring mature gametophytes from any sexual or apomictic *Boechera* accession sampled in this study. (B) Schematic drawing of developmental stages and reproductive tissues sampled.

**Table 1:** *Boechera* accessions used for transcriptional profiling in this study and reproductive tissues sampled.

| ***Boechera* species** | **accession number** | **ploidy** | **reproductive mode** | **reference** | **tissues sampled** |
| --- | --- | --- | --- | --- | --- |
| *B. holboellii* | ES786 | 2x | sexual | (Mau et al., 2015, Aliyu et al., 2010) | mature ovules |
| *B. stricta* | LTM | 2x | sexual | (Lee et al., 2017, Schranz et al., 2005) | nucellus MMC, nucellus FMS, mature ovules |
| *B. williamsii* | B12-558 | 2x | sexual | (Mau et al., 2015) | nucellus MMC, mature ovule |
| *B. williamsii* | B12-1524 | 2x | apomictic | (Mau et al., 2015) | nucellus MMC |
| *B. divaricarpa* | ES517 | 2x | apomictic | (Mau et al., 2015, Aliyu et al., 2010) | nucellus MMC, mature ovule |
| *B. pallidifolia* | B12-1578 | 2x | apomictic | (Mau et al., 2015) | nucellus MMC, nucellus uninucleate gametophyte, mature ovules |
| *B. pallidifolia* | B12-1599 | 3x | apomictic | (Mau et al., 2015) | nucellus MMC, nucellus uninucleate gametophyte, mature ovules |

All accessions have previously been described as indicated under “reference”. Accessions represent different reproductive modes and ploidy levels.

Subsequent to preprocessing of data by quality control and trimming, reads were mapped against the *B. stricta* reference genome (Lee et al., 2017). After counting of unique reads mapped to exonic regions, 15‘577 and 14‘717 genes expressed (≥ 10 TMM normalized read counts (Tarazona et al., 2015)) were identified in nucellus tissues harbouring an AIC/MMC, and 16‘481 and 19‘437 at onset of gametogenesis, in addition to 18‘211 and 17‘461 genes active in mature ovules, in any sample from apomictic or any sample from sexual accessions, respectively (Figure 1). Furthermore, 12‘051 genes shared expression in all samples.

First, we were interested in changes of transcriptional regulation between different developmental stages within each reproductive mode. Statistical analysis identified 7‘004 genes as significantly differentially expressed (FDR≤0.05 after Benjamini Hochberg adjustment) when contrasting the developmental stages of MMC, FMS, and mature ovules in all sexual accessions using a quasi-likelihood F-test in an ANOVA-like analysis (Table S2A) (Robinson et al., 2009). To gain insights into the molecular functions enriched in this set of genes, we made use of the annotation of *A. thaliana* homologues in the *B. stricta* genome and the gene ontology (GO) annotations for them. In total, GO analysis identified terms of 17 molecular functions to be significantly enriched (p < 0.01), including „translation initiation factor activity“, „structural constituent of ribosome“, „RNA binding“, and „helicase activity” (Table 2). Furthermore, activity of RNA polymerase II, metabolic pathways, and functions related to transport and ubiquitinylation were identified, in addition to chromatin related functions (i.e. „chromatin binding“, and „histone binding” as nearly significant with p = 0.01). Similarly in apomicts, analysis identified 9’136 significantly differentially expressed genes (DEGs) (Table S2B). From this set of genes, 34 GO terms of molecular functions were identified as significantly enriched (p < 0.01) (Table S3). While the top terms were shared between sexual and apomictic reproduction, suggesting a general role for reproductive development, additional categories pointed towards the importance of signaling pathways and redox regulation.

**Table 2:** Molecular functions identified as enriched in a gene ontology analysis on the 7‘004 genes identified as DEGs at different stages of development in all sexual accessions.

| GO.ID | Term | Annotated | Significant | Expected | p Value |
| --- | --- | --- | --- | --- | --- |
| GO:0003735 | structural constituent of ribosome | 221 | 152 | 67.52 | < 1e-30 |
| GO:0003723 | RNA binding | 1064 | 458 | 325.06 | 1.20E-16 |
| GO:0004386 | helicase activity | 148 | 69 | 45.22 | 3.80E-05 |
| GO:0004427 | inorganic diphosphatase activity | 12 | 10 | 3.67 | 0.00024 |
| GO:0003743 | translation initiation factor activity | 80 | 39 | 24.44 | 0.00048 |
| GO:0008320 | protein transmembrane transporter activity | 41 | 23 | 12.53 | 0.00059 |
| GO:0003843 | 1,3-beta-D-glucan synthase activity | 11 | 9 | 3.36 | 0.00067 |
| GO:0046933 | proton-transporting ATP synthase activity | 19 | 13 | 5.8 | 0.00075 |
| GO:0001055 | RNA polymerase II activity | 17 | 12 | 5.19 | 0.00078 |
| GO:0004175 | endopeptidase activity | 298 | 118 | 91.04 | 0.00110 |
| GO:0044877 | protein-containing complex binding | 167 | 74 | 51.02 | 0.00129 |
| GO:0003779 | actin binding | 92 | 42 | 28.11 | 0.00159 |
| GO:0003682 | chromatin binding | 132 | 55 | 40.33 | 0.00431 |
| GO:0005198 | structural molecule activity | 320 | 194 | 97.76 | 0.00605 |
| GO:0016831 | carboxy-lyase activity | 60 | 28 | 18.33 | 0.00622 |
| GO:0050897 | cobalt ion binding | 50 | 24 | 15.28 | 0.00711 |
| GO:0061650 | ubiquitin-like protein conjugating enzyme activity | 43 | 21 | 13.14 | 0.00900 |
| GO:0042393 | histone binding | 78 | 34 | 23.83 | 0.01004 |

To study differences in regulation in more detail, we in addition performed a pairwise comparison on all samples from nucelli at MMC stage *versus* all samples from mature ovules of sexual accessions. Thereby, 7‘928 DEGs were identified, 4‘219 higher expressed in nucellus tissues and 3‘709 in mature ovules (Figure 2, Table S4). Consistently, 6‘140 DEGs were identified both in the pairwise comparison of the two and the ANOVA-like analysis of all three developmental stages. Analysis of enriched biological processes based on the genes upregulated either at MMC-stage or in mature ovules further supported evidence for the importance of ribosomal pathways and translational control during megasporogenesis (Table S5A), while for mature ovules enriched processes included such related to hormonal pathways, epigenetic regulatory pathways, signaling, development and transcriptional regulation (Table S5B).

**Figure 2:**
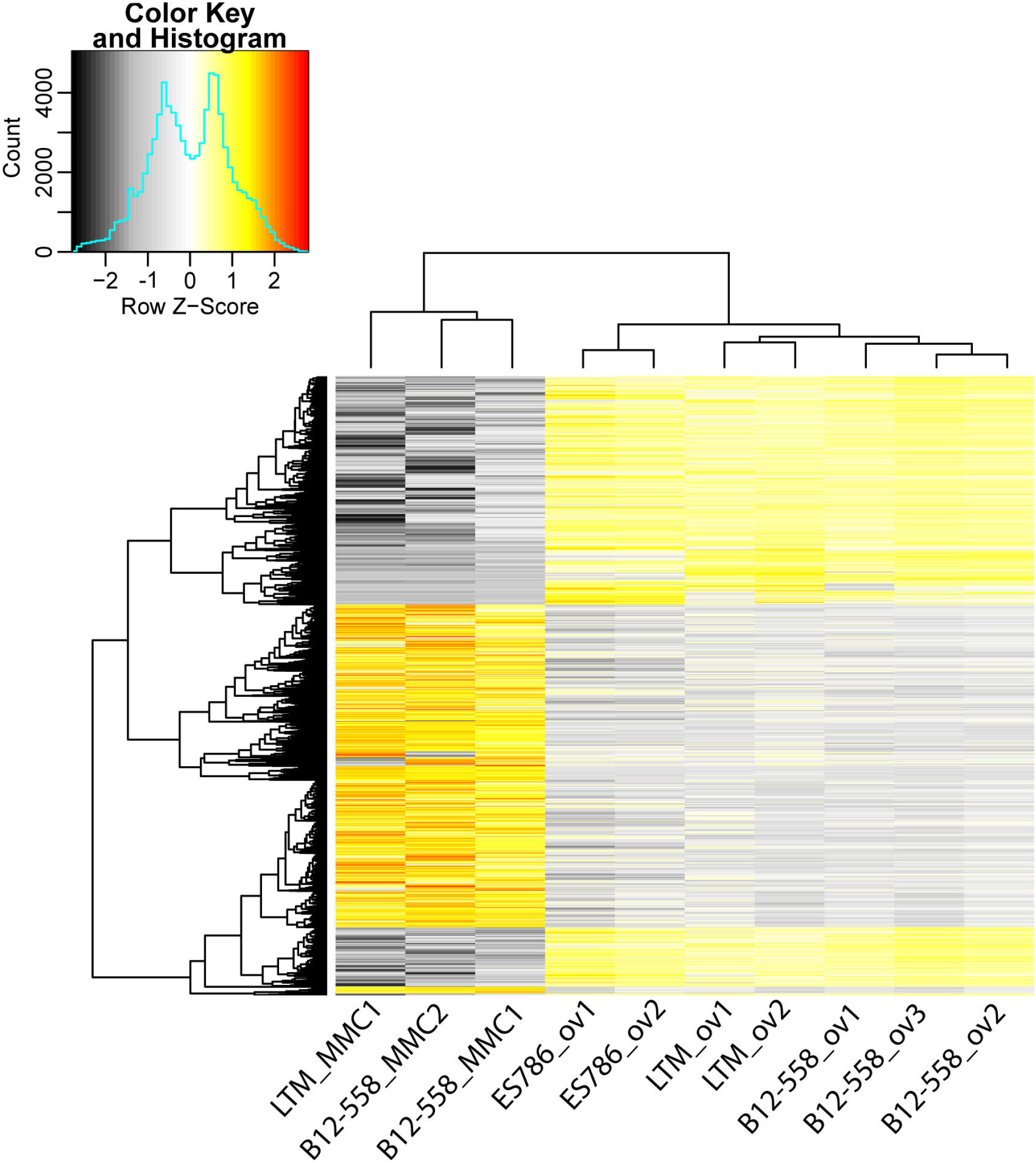
Heatmap of normalized read counts. (A) 7928 genes differentially expressed in nucellus tissues harbouring the MMC and mature ovules in sexual *Boechera* based on log2-scale TMM-normalized read counts. (B) Hierarchical clustering of genes and samples was based on euclidian distance and hierarchical agglomerative clustering. Colors are scaled per row and red denotes high expression and black denotes low expression.

### Differential expression analysis suggests importance of RNA helicases for reproduction

In agreement with their supposed functional importance, gene ontology analysis identified 69 DEGs corresponding to 71 *B. stricta* homologues annotated to the molecular function of “helicase activity” (Table S6). Furthermore, 75 RNA helicases were identified to be comprised in the 7‘004 DEGs when searching for homologues in *A. thaliana* based on annotations of the *B. stricta* genome (Figure S1) (Xu et al., 2013, Lee et al., 2017). Interestingly, for *A. thaliana ESP3* two homologues (*Bostr*.*3359s0111, Bostr*.*7867s0308*) were amongst the DEGs, in addition to two homologues of *FAS4* (*Bostr*.*17016s0021, Bostr*.*17016s0022*, hereafter referred to as *0021* and *0022*). Despite significant differences in expression levels, *0022* is only low or moderately expressed in the tissues analysed. Furthermore, DEGs comprised homologues of more than half of the *A. thaliana* DEAD-box RNA helicases, including *RH17, RH22, RH29, RH36*, and *RH39* (Table S6). Apart from genes annotated as homologues of *ESP3* (Herr et al., 2006), additional RNA helicases involved in epigenetic regulatory pathways are represented, including homologues of *DICER LIKE1* (*DCL1*), *DCL2, DCL4, SILENCING DEFECTIVE3, DEFECTIVE IN RNA-DIRECTED DNA METHYLATION 1, CHR11*, and of different genes involved in regulation of genome stability (Table S6). Furthermore, *VASA-LIKE* was identified in this analysis, which in *A. thaliana* is predominantly expressed during megasporogenesis and important for reproductive development (Schmidt et al., 2011, Kiefer et al., 2020).

To date, little is known about the gene regulatory pathways controlling aspects of specification of cells of the mature gametophyte in apomicts. For pseudogamous apomicts, it can be envisioned that regulatory mechanisms are in place to safeguard the single fertilization process and to sustain suppression of embryogenesis in its absence. To identify potential candidates in these processes we investigated differential expression in ovule tissues from sexual and apomictic accessions. We identified a total of 917 DEGs, 519 of which were expressed at significantly higher levels in sexual accessions (Table S7, Figure S2).

Previous cell type-specific transcriptional analysis of *A. thaliana* egg, central cell, and synergids have identified genes predominantly expressed in these cell-types, which are of putative importance for their specification, identity, and function (Wuest et al., 2010). To uncover such genes enriched in sexual gametophytes but changed in regulation in apomicts, we were particularly interested in genes expressed at significantly higher levels in ovules from sexual as compared to apomictic *Boechera*. Amongst the candidates fulfilling this criterion, 12 *Boechera* homologues of 11 *A. thaliana* genes were identified (Figure S3), interestingly including *FAS4* (*0021*) and *CHR34*. In addition, DEGs included homologues of additional egg cell preferentially expressed genes. These are *At4G36150*, encoding for a disease resistance protein of the TIR-NBS-LRR class family, and *JMJC DOMAIN-CONTAINING PROTEIN 27 (JMJ27)*. In addition, of the genes enriched in other cells or any cell of the mature *A. thaliana* gametophyte (Wuest et al., 2010), including *KIP-RELATED PROTEIN 7* (*KRP7*), homologues of 11 *A. thaliana* genes were significantly higher expressed in ovules of apomictic as compared to sexual accessions, suggesting their deregulation (Figure S3).

Importantly, our study supports previous evidence that several RNA helicases are crucial for megasporogenesis. Nevertheless, in mature ovules of sexual as compared to apomictic accessions, 5 RNA helicases were identified as DEGs (Figure S3). Like *0021* and *CHR34*, also a homologue of *RH1*, and Bostr.*30275s0126*, annotated as *ESP3* homologue, were expressed at higher levels in sexual accessions. In contrast, a homologue of *ROCK-N-ROLLERS* (*RCK*) was significantly higher expressed in mature ovules of apomictic as compared to sexual accessions. In summary, we identified new genes and especially additional RNA helicases as candidates for sexual and apomictic reproduction.

### Evolution of *ESP3, MEE29* and closely related genes might shape reproductive development

Transcriptional analysis identified distinctive regulation of different homologues of *ESP3*. Therefore, we got interested in their evolutionary history. To this aim, we searched genomes of six selected Brassicaceae for homologues (Table S8, Figure S4). A blast search suggested a close relation of *ESP3* and *MEE29*, and we further identified *Bostr*.*3359s0111* and *Bostr*.*23794s0568* as positional orthologues of *AtESP3* and *AtMEE29*, respectively. In addition, *Bostr*.*30275s0126* was identified as orthologue of the uncharacterized gene *At4G16680*. Another locus, *Bostr*.*7867s0308*, encoding a helicase was identified in *B. stricta*. Its closes relative in *A. thaliana* was *At1g32490*, which is *ESP3*. No orthologue of *Bostr*.*7867s0308* was found in the genomes of *Euclidium syriacum, Eutrema salsugineum*, and *A. thaliana*, while syntenic orthologues could be identified in *Arabidopsis lyrata* and *Arabis alpina*, suggesting independent events of loss in *Euclidium syriacum, Eutrema salsugineum*, and *A. thaliana*. A phylogenetic maximum likelihood analysis by raxml-ng (Stamatakis, 2006) further suggests that all genes likely originate from a common ancestor, but *ESP3* diverged early form all other genes (Figure S5). Furthermore, *At4G16680* and homologues form a clearly distinctive group, and homologues of *MEE29* are closer related to *Bostr*.*7867s0308*. A dynamic evolutionary diversification is also reflected by a high degree of sequence similarity in exonic regions and lower degrees of sequence similarities in non-coding intronic or flanking regions of orthologues from the different species (Figure S6). In particular for *Bostr*.*7867s0308*, sequence similarities were largely restricted to exonic regions suggesting changes in regulation (Figure S6D).

### One orthologue and two paralogues of *A. thaliana FAS4* are present in *Boechera*

Based on expression patterns of *FAS4* homologues, we hypothesized functional diversification of the variants with potential relevance for gametogenesis. To study their evolutionary relation, we searched genomes of the six Brassicaceae species for homologues of *AtFAS4* (*At1G33390*) and its positional neighbors, *At1G33360* and *At1G33400*. Sequences showing high homology to *AtFAS4* could be identified in all species considered (Figure 3A, 3B, Table S8). However, while *At1g33360* and *At1g33400* were syntenic (with additional genes inserted between the two loci) in all taxa included in the comparison, sequences with high homology to *FAS4* were found to be highly variable in terms of genomic localization. In *A. alpina, E. salsugineum* and *E. syriacum* these sequences were located on other scaffolds or chromosomes than the two loci which are neighbouring in *A. thaliana*. Only in Brassicaceae taxa belonging to evolutionary lineage I (*Arabidopsis, Boechera*) (Walden et al., 2020), the sequences homologous to *FAS4* were found to be located between sequences orthologous to the *A. thaliana* upstream and downstream genes used in the blast search. Hence, the setup found in *A. thaliana* evolved in lineage I only. Besides in *B. stricta* sequences orthologous to *FAS4* had also been multiplied in *A. lyrata*. In *B. stricta* one homologous sequence was found in synteny with the *A. thaliana* upstream and downstream genes on scaffold 3359 (gene *0032*). The two additional sequences were found to be located in tandem on scaffold 17016 (genes *0021, 0022*). These putatively represent *Boechera* specific paralogues likely derived from the original gene. A common origin is suggested by a high sequence similarity of the coding regions and proximal promoter as shown in the VISTA plot (Figure 3B,C), also showing two short conserved regions several kb upstream of the coding region.

**Figure 3:**
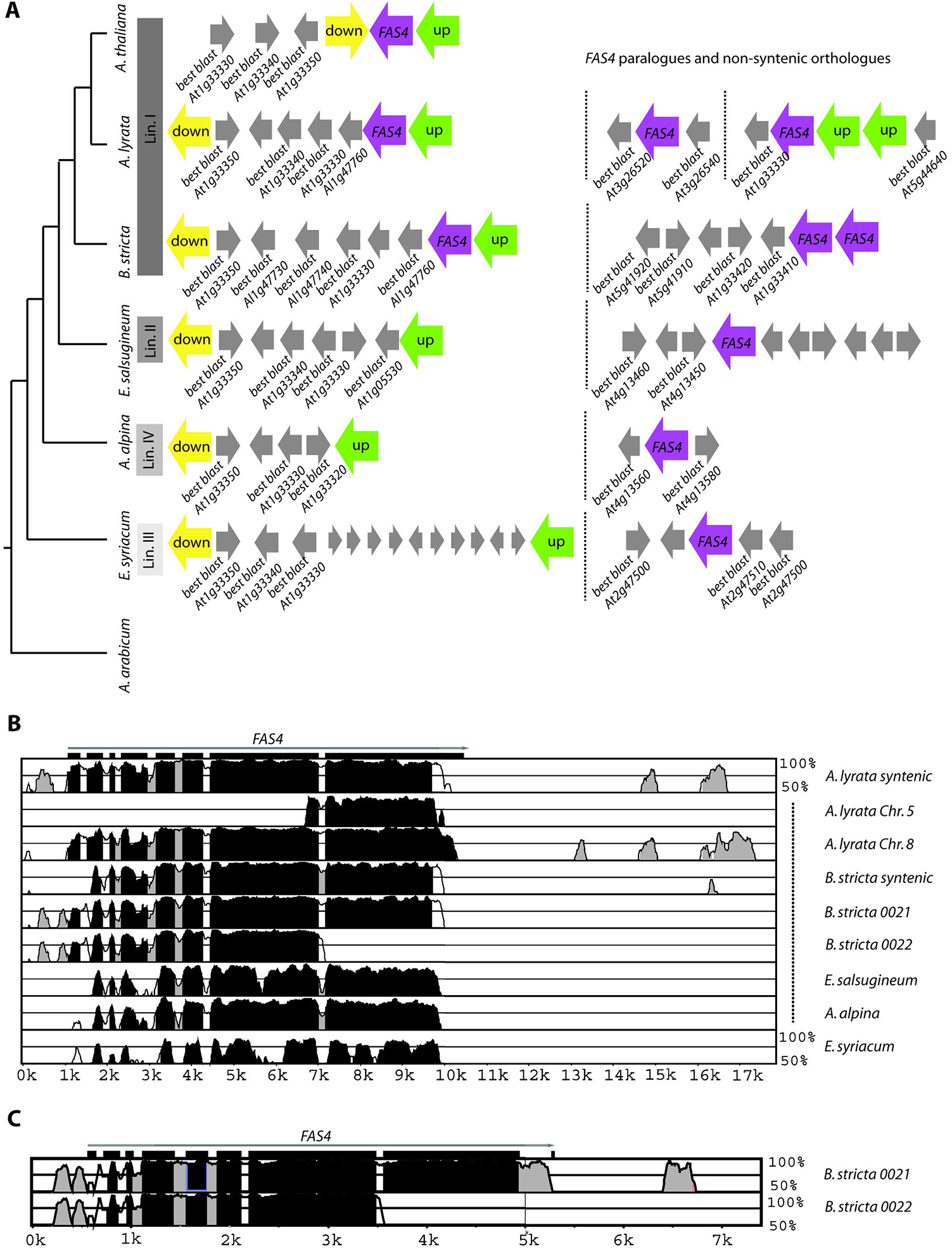
Evolutionary analysis on *FAS4*. (A) Schematic representation of relatedness of analysed Brassicaceae species drawn according to (Kiefer et al., 2019). Gene arrangements of *FAS4* (purple) orthologues (on the left side) and paralogues and non-syntenic orthologues (on the right side) as well as upstream (yellow) and downstream (green) genes. Best blast hits of neighboring genes are given when *FAS4* orthologue is not syntenic. (B) mVISTA plot using the genic and intergenic sequence of *FAS4* genes from all six Brassicaceae species using *A. thaliana* as base. X-axis is alignment position, y-axis is percent identity. Genic sequences were shown to be highly similar. In addition, one intergenic segment was found to be conserved except for *E. salsugineum, A. alpina* and *E. syriacum*. (C) mVISTA plot between *0021* and *0022* shows high sequence conservation in proximal regulatory regions upstream and exons 2-7, whereas the 8^th^ exon is lacking in *0022*.

In *A. lyrata* the situation differed slightly. One sequence was also found to be syntenic compared to the gene order in *A. thaliana* (LG1; *Al1g47770*, neighbours are *Al1g47710* and *Al1g47780*; five genes interspersed between *Al1g47710* and *Al1g47770*). However, two additional sequences were found on LG8 (*Al8g14950*; neighbours *Al8g14970* and *Al8g14970* show high homology to *At1g33340*, which is the upstream gene of *FAS4*) and LG5 (*Al5g15310*). Blast of the neighbouring genes of the sequences on scaffold 17016 in *B. stricta* and on LG8 and LG5 in *A. lyrata* to the *A. thaliana* reference genome do not allow to draw any conclusion on a common duplication event in *A. lyrata* and *B. stricta* leading to the multiple sequences showing homology to *FAS4*. Furthermore, upstream regulatory regions and N-terminal regions did not show evident similarities for *AL5g15310* as compared to any other of the homologues from Brassicaceae (Figure 3B). In addition, phylogenetic analysis rather indicates a more complex evolutionary history and supports a likely origin of the paralogues by a later duplication event in *Boechera* (Figure S7). Together this suggests that the evolution of *FAS4* and its homologues is marked by high mobility in the Brassicaceae genomes, potentially having consequences on regulation and function.

### Phylogenetic reconstruction of *FAS4* positional orthologues and paralogues identifies distinct clusters

To gain insights into aspects of the evolution of RNA helicases in *Boechera*, previously a target enrichment study has been performed on a number of *Boechera* accessions (Kiefer et al., 2020). Mapping the sequencing reads that have been generated in this study revealed that due to their high similarity sequences were captured covering large parts of both loci including also non-coding regions upstream and downstream of the genes. Phased alleles of *0032* could be obtained for nine sexual, six facultative and four apomictic accessions and phased alleles for *0021* were obtained for six sexual, two facultative and two apomictic accession (Table 3, Figure 4). In *0032*, premature stop codons were identified in two facultative (ES911; B12-1591) and one sexual accession (ES820). Premature stop codons occurred in both alleles in the individuals but in case of the facultative apomicts not at the same position. In *0021* premature stop codons were identified in the two facultative apomicts which also showed premature stop codons in *0032*.

**Figure 4:**
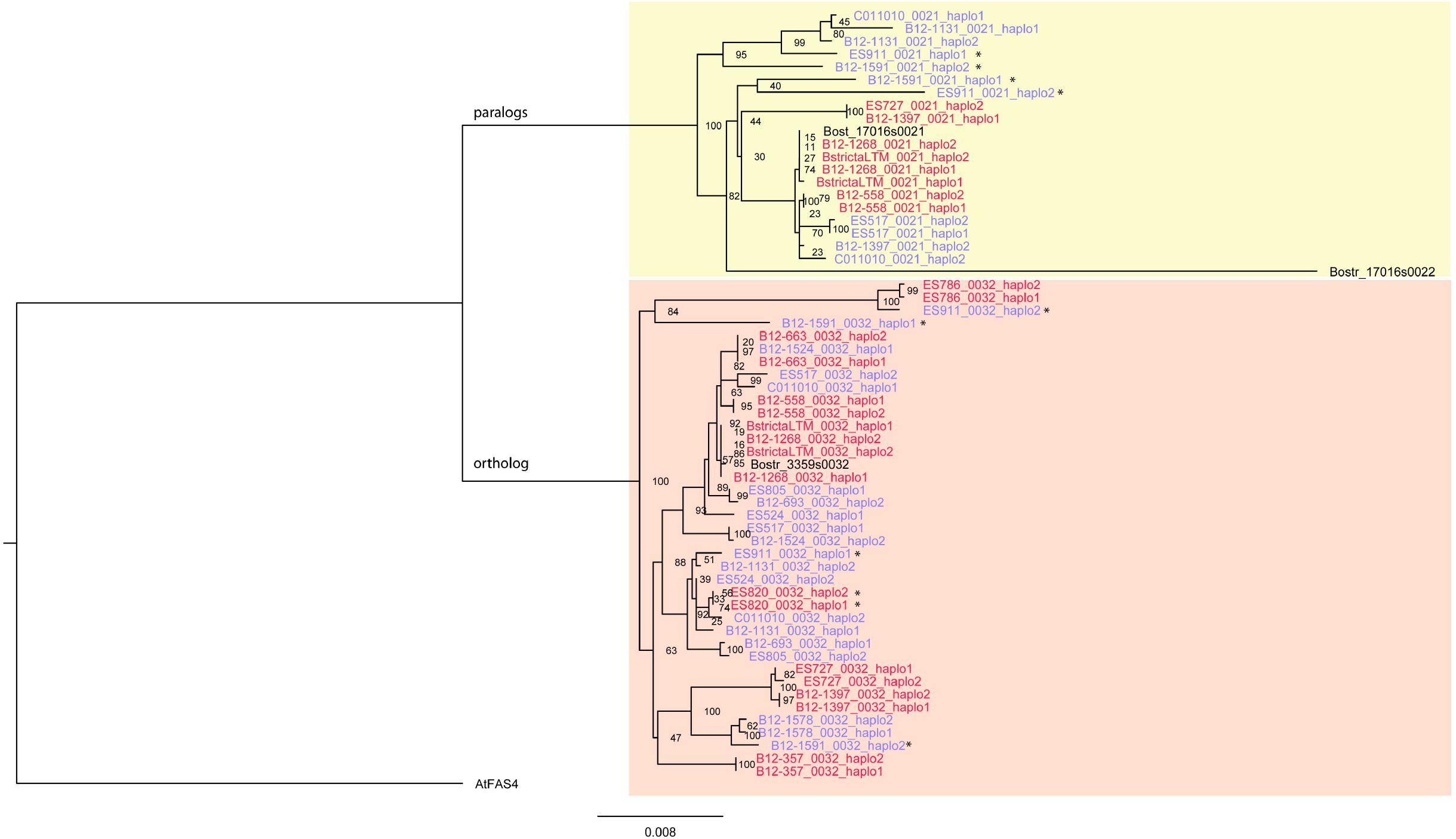
Phylogenetic analysis. This was done by raxml-ng using the HKY+G model and an alignment comprising all identified phased alleles of *0021* and *0032*, as well as the sequences of *0021, 0022* and *0032* from the *B. stricta* draft genome and *FAS4* from *A. thaliana* as outgroup. *0032* and *0021/0022* form sister groups. While in the subclade comprising *0032* alleles apomicts are more interspersed, they tend to cluster in the clade containing *0021*. It should be noted that apomicts also often contain distantly related alleles reflecting their hybrid nature. Alleles from apomictic accessions are marked in blue, alleles from sexual accessions in red. Alleles harbouring a stop codon or frame shift mutation are indicated with a star.

**Table 3:**
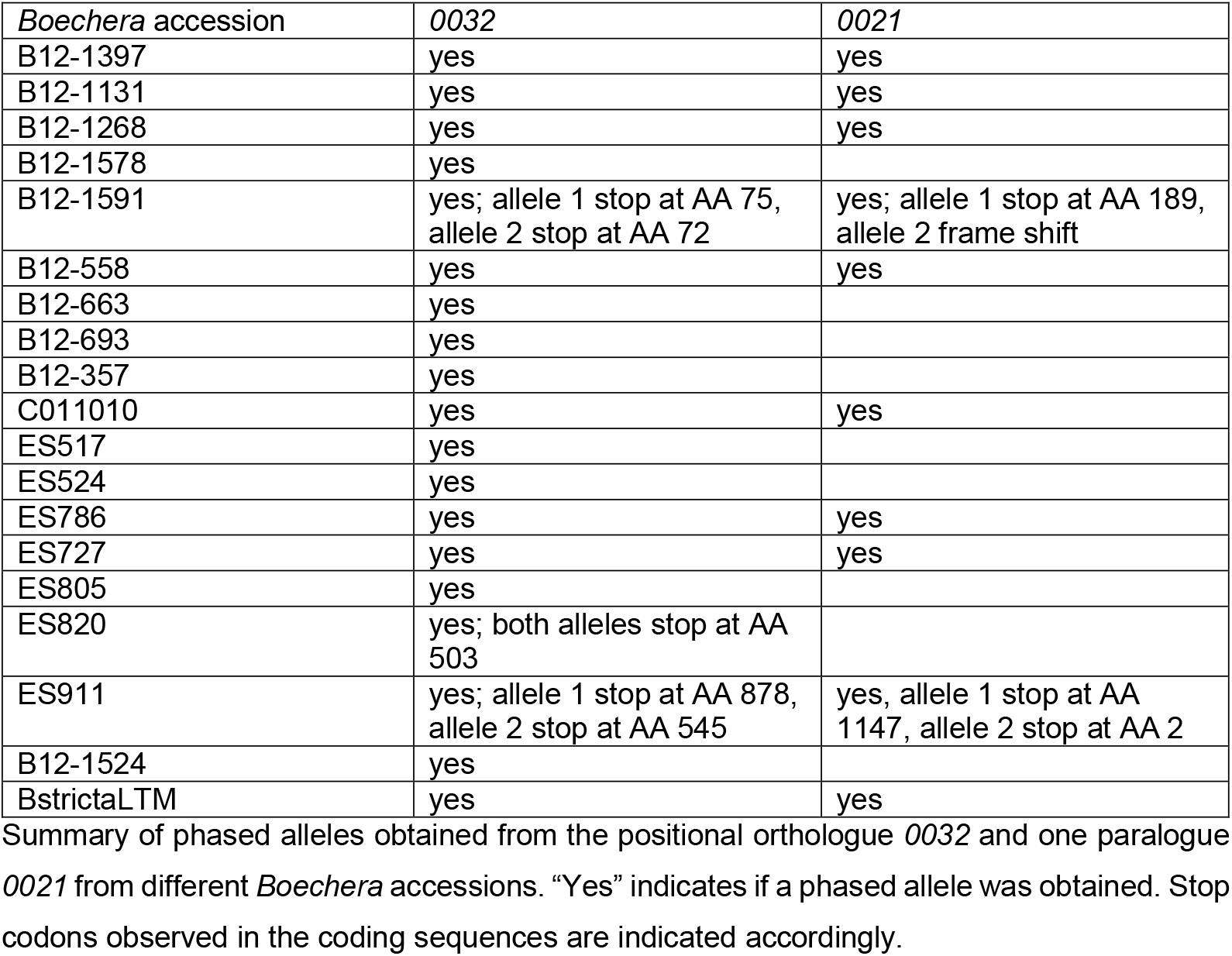
Summary of alleles.

Phylogenetic reconstruction based on all CDS of *0032, 0021* and *0022* using *AtFAS4* as outgroup clearly split the genes into two groups – one group contained all *0032* sequences while the other group contained *0021* and *0022* (Figure 4). In the clade containing the *0032* sequences alleles harbouring premature stop codons occurred in different subclades. This suggests that non-functionalization of alleles of *0032* happened multiple times independently. The same is true for *0021*, as also here alleles containing premature stop codons occur in different subclades.

While for *0032* alleles originating from apomicts and facultative apomicts do not cluster but are found in all subclades, alleles of 0021 originating from apomicts or facultative apomicts do cluster in one group, except for the two alleles found in ES517 and one allele found in C011010. Further, most sexual accessions contain alleles, which are closely related while (facultative) apomictic accessions harbour quite divergent alleles often found in different subclades. This underlines the apomictic nature of *Boechera* apomicts.

### Mutant alleles of *A. thaliana FAS4* lead to segregation distortion

The characteristic expression pattern in reproductive tissues and the evolutionary diversification suggests potential roles of *FAS4* for gametogenesis. In line with this, we observed localization of *At*FAS4-PmTurqouise fusion proteins in female gametophytes, and of *At*FAS4-PmTurquoise and *Bs*FAS4-mVenus in anther tissues (Figure S8, Supporting information). To investigate the potential roles of *FAS4* for plant reproductive development, we studied independent *A. thaliana* lines carrying mutant alleles, which are a line carrying a T-DNA insertion in the 4^th^ intron (*fas4-1*), and three lines generated by CRISPR/Cas9 harbouring mutations in the 4^th^ exon of *FAS4*, hereafter referred to as *fas4-2, fas4-3*, and *fas4-4* (Supporting Information, Table S9). The lines showed defects in fertility with aborted or unfertilized ovules at 44.7% (n = 322) in *fas4-1/FAS4*, 51.3% (n = 156) in *fas4-2/FAS4*, 40.2% (n = 162) in *fas4-3/FAS4*, and 42.9% (n = 224) in *fas4-4/FAS4*, as compared to 5.2% in the wild-type (n = 532), pointing towards a gametophytic defect.

In agreement with the frequencies of abortive ovule development, severe segregation distortion was observed in offspring of heterozygous mother plants when scoring the ratio of phosphinothricin resistant to sensitive seedlings in *fas4-1/FAS4*, with only 13.1% (n = 808) of resistant seedlings recovered. This suggests defects in female and male gametogenesis. In agreement, transmission efficiencies through both parents were drastically reduced after reciprocal crosses to wild-type (Table 4).

**Table 4:**
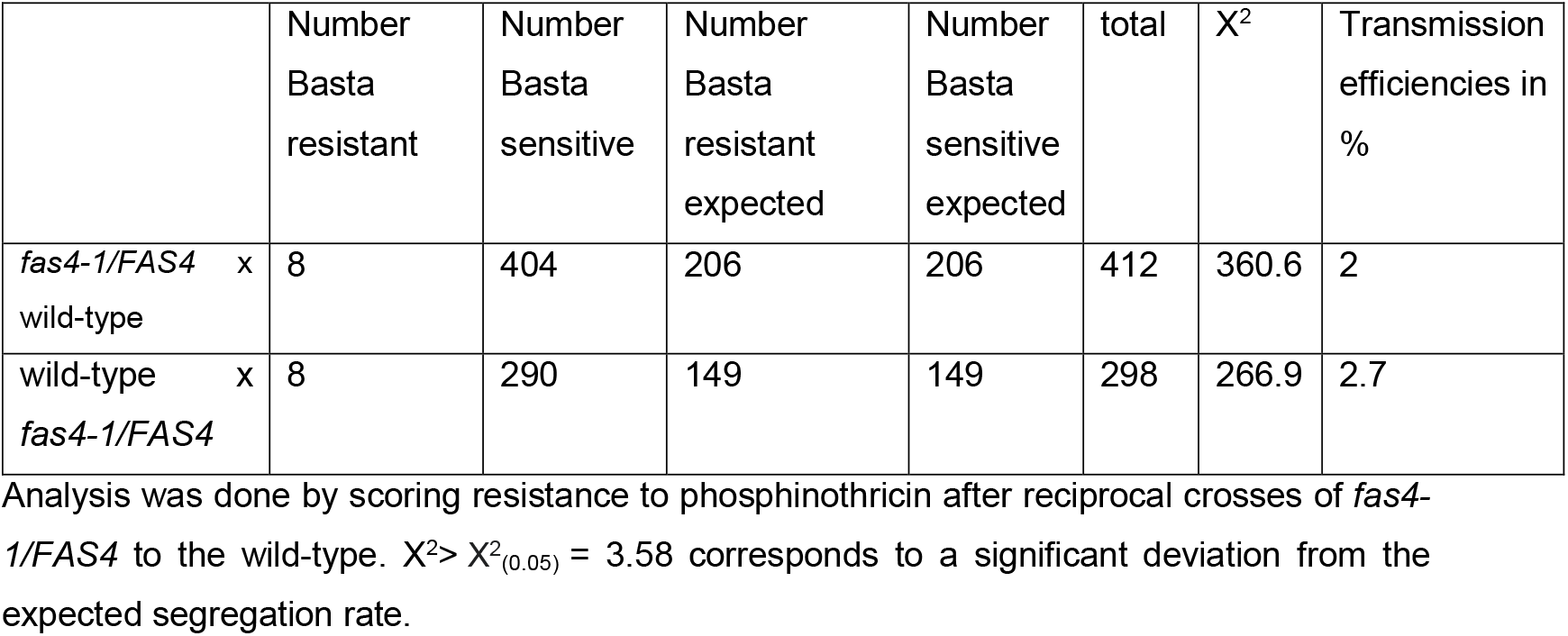
Transmission efficiencies of mutant *fas4-1* allele.

### *FAS4* is crucial for gametogenesis in *A. thaliana*

To study pollen development, we first applied Alexander Staining to identify the proportion of viable pollen grains (Ross et al., 2010). For *fas4-1/FAS4*, we identified 70% of dark pink viable pollen, 14% of aborted pollen (light blue), and 16% of light pink pollen (n = 1024) (Figure 5A), as compared to 91% of viable and 9% of aborted pollen in wild-type (n = 487), where the category of light pink pollen was not represented. This suggests that the light pink pollen might have developmental defects. To gain more precise insights into male gametogenesis we further investigated pollen from freshly opened flowers by DAPI staining. We observed 63% of normally developed tricellular pollen in addition to different developmental defects, including 4% of aborted microspores, 20% of uninucleate or binucleate pollen, suggesting defects of PMI or PMII, and 13% with decondensed chromatin of the vegetative nucleus (n = 227) (Figure 5B). In contrast, 91% of mature tricellular pollen in addition to 9% bicellular pollen (n = 154) was observed in wild-type (Figure 5C).

**Figure 5:**
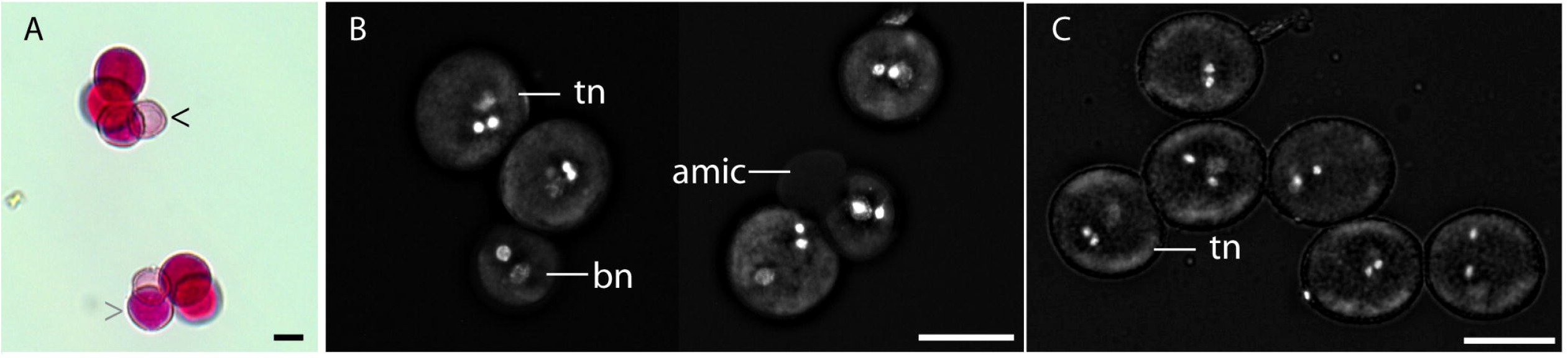
Pollen development and viability. (A) Alexander Stain (Ross et al., 2010) was used to test for pollen viability in *fas4-1/FAS4*. Viable pollen showed dark pink staining, while aborted pollen showed a light blue color (black arrow). In addition, a category of light pink pollen (grey arrow) was observed. (B,C) Epifluorescence microscopy on pollen of freshly opened flowers stained with DAPI identifies trinucleate pollen (tn), binucleate pollen (bn), and aborted microspores (amic) in *fas4-1/FAS4* (B), and trinucleate pollen in wild-type (C). Scale bars are 20 µm.

Similar to mitotic defects during male gametogenesis we observed delayed or aborted progression of female gametophyte development (Figure 6, Figures S9, S10). In *fas4-1/FAS4* at 2 days after emasculation (2 DAE) we observed 44.3% of mature wild-type gametophytes (n = 467), and an additional 3.9% showed mature gametophytes but with unfused polar nuclei. Furthermore, 45.6% of ovules harboured gametophytes showing delayed or aborted progression through gametogenesis, or untypical numbers or positioning of gametophytic nuclei (Figure S9). And 6.2% of ovules were either collapsed or comprised gametophytes showing developmental arrest at onset of gametogenesis. Similarly, also in wild-type we observed unfused polar nuclei in 5.3% of gametophytes (n = 131) and 7.7% of collapsed ovules or ovules harbouring gametophytes that were arrested early during gametogenesis. Nevertheless, 87.0% of gametophytes were fully matured (Figure S9). In agreement, based on the T2 generation before backcrossing to the wild-type (Supporting Information), clearing of pistils around anthesis identified 74% normally developed gametophytes and 26% delayed or aborted gametophytes in *fas4-2/FAS4* (n = 19). Similarly, young developing siliques of *fas4-3/FAS4* showed 57% delayed or arrested gametophytes, and 42% mature gametophytes or young developing seeds (n = 14), while in *fas4-4/FAS4* 48% of wild-type like mature gametophytes or young seeds were observed (n = 23), whereas the remainder of the ovules harboured arrested gametophytes or untypical numbers or positioning of gametophytic nuclei (Figure 6). Even though similar phenotypes were observed in all three lines, numbers of observations differed. This might be due to the rather low numbers of ovules observed for the CRISPR/CAS9 generated lines.

**Figure 6:**
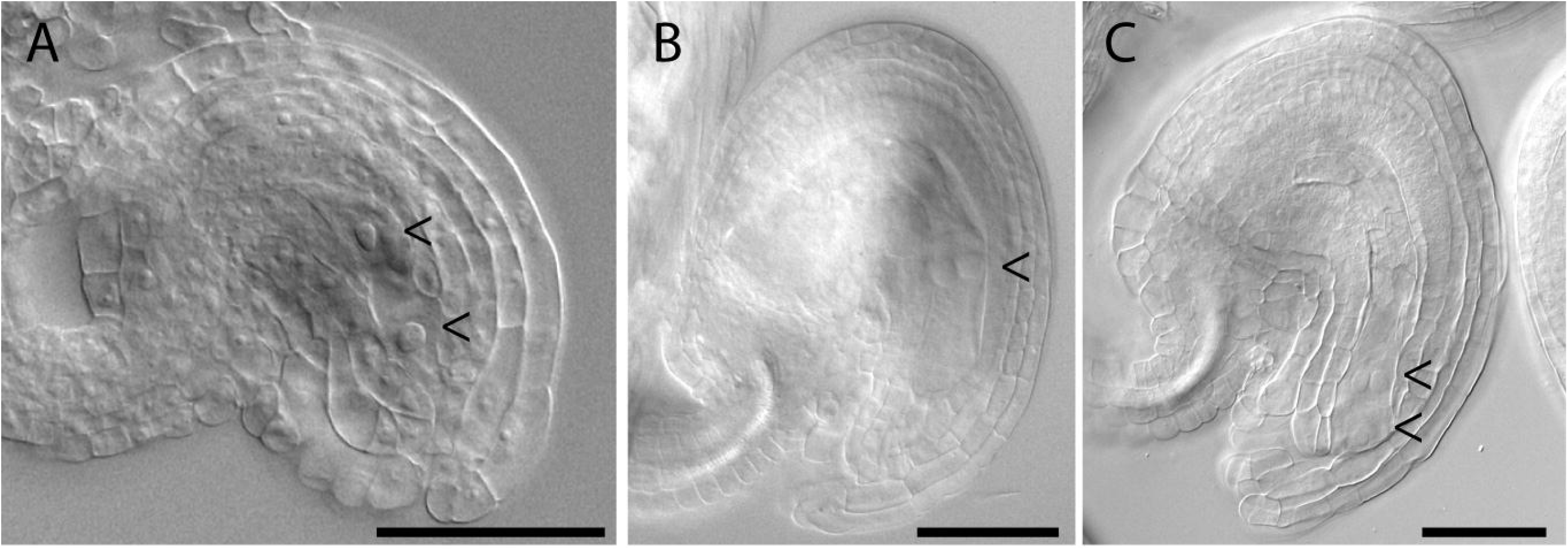
Analysis of female gametogenesis. (A) Two-nucleate wild-type like gametophyte in *fas4-1/FAS4*. (B) Gametophyte with two nuclei in odd positions in *fas4-4/FAS4*. (C) Gametophyte with three nuclei in *fas4-1/FAS4*. Gametophytic nuclei are indicated with arrows. Scale bars are 50 µm.

To elucidate in more detail when developmental defects were first visible, we made use of *fas4-1/FAS4* plants harbouring the p*AKV::H2B-YFP* construct labelling gametophytic nuclei (Rotman et al., 2005). We scored frequencies of developmental alterations in buds of different stages (Figure S10A-E, Table S10). Therefore, we broadly categorized buds based on the developmentally most advanced gametophytes observed typically constituting of wild-type. We classified stages before first and second mitotic division (1-2 gametophytic nuclei), after second mitotic division (up to 4 gametophytic nuclei), and after third mitotic division (8 or 7 gametophytic nuclei). This suggested that formation of the FMS was typically normal and developmental defects accumulated during mitotic division (Figure S10A-E, Table S10).

To gain first insights into the question, if potential mitotic defects might lead to altered ploidy in gametophytic nuclei, we introgressed the p*WOX2::CENH3-GFP* marker line into *fas4-1/FAS4* to visualize individual chromosomes (De Storme et al., 2016). While no clear indication was observed for higher ploidy of individual gametophytic nuclei (Figure 10F-J), evidence was found for potential alterations in planes during mitotic divisions (Figure S10H), alterations in migration (Figure S10I), or separation of gametophytic nuclei (Figure S10J), which were observed in 22% of the gametophytes (n = 167). In contrast in the p*WOX2::CENH3-GFP* line serving as wild-type control, similar defects were only observed in 3% of gametophytes (n = 308). Together, this demonstrated that *At*FAS4 is crucial for male and female gametogenesis and potentially for mitotic divisions during gametogenesis. Interestingly, this importance is likely based on roles associated to ribosome biogenesis (Supporting Information, Figure S11).

## Discussion

A striking feature of gametogenesis in higher plants is that developmental gradients along polar cell axis and asymmetric divisions play important roles for cell fate determination (Tekleyohans et al., 2017). In this study, we identified crucial roles of *AtFAS4* for gametogenesis. In female gametophytes of lines carrying mutant *FAS4* alleles, aberrations in separation, positioning, and numbers of gametophytic nuclei were observed, suggesting mitotic defects and potentially alterations in polar development. To establish and sustain polarity in the syncytial gametophyte, localized RNA deposition and processing might be involved. To mediate spatiotemporal regulation of translation, RNA helicases and other RNA binding proteins associated to RNPs supposedly are major components. While from investigations of *Drosophila* oogenesis and early embryogenesis the relevance of an interplay of different RNA binding proteins for polar development has been uncovered (Bansal et al., 2020, Hughes and Simmonds, 2019), to date little is known about the molecular machineries modulating the translatome during plant reproduction. Nevertheless, for tobacco pollen large cytoskeleton associated RNPs have been described (Honys et al., 2009). Here, the molecular machinery contains rRNAs and mRNAs, which remain translationally silent during pollen development and only get activated during pollen tube growth and sperm cell delivery to the female gametophyte (Honys et al., 2009). In contrast, little is known about the roles of RNA helicases associated to ribosomes in specification and function of the female gametes.

Based on the predominant expression of *AtFAS4* in synergids, it was identified to be a promising candidate for gametogenesis. Indeed, it is a unique gene in *A. thaliana* and required for male and female gametogenesis. On the female side, defects conferred by mutant *FAS4* alleles partially resemble phenotypes induced by a mutant allele of *SWA3* (Huang et al., 2010, Liu et al., 2010). It is thus possible that FAS4 and SWA3 are components of the same ribosome associated pathway and control aspects of mitotic progression during gametogenesis. However, while localization of *At*FAS4 in maturing and mature female gametophytes could be confirmed, on the male side both *A. thaliana* and *B. stricta* orthologues are present in anther tissues surrounding pollen. This likely affects pollen maturation and viability, and might impact pollen germination or pollen tube growth not investigated here. Such effects in combination could explain that mutant alleles are only transmitted at minor frequencies through sperm, although a higher percentage of pollen carrying a mutant allele matures.

Interestingly, genes involved in ribosome biogenesis are also important for fertilization. Apart from developmental defects during gametogenesis, plants carrying a mutant allele of *MAA3* display problems in pollen tube attraction (Shimizu et al., 2008). Successful pollen tube attraction and double fertilization depend on the specification of the synergids, but also on small cysteine rich proteins exocytosed from the egg cell to activate the sperm cells and to prevent delivery of multiple sperms (Sprunck et al., 2012). The regulatory system underlying is complex and mediates a balance between conserved developmental processes and alternations from these programs to create evolutionary opportunities. This is for example demonstrated by the recent description of mechanisms leading to polyploidization by specific fertilization of egg cells only with additional sperms, unlike central cells. This prevents aberrations from the maternal to paternal genome ratio critical for endosperm development (Mao et al., 2020). The delicate balance of male female interaction to allow pollen tube attraction, discharge, and sperm activation supposedly gets more complex in pseudogamous apomicts, where two sperms are delivered but only a single fertilization event of the central cell typically takes place. It can be hypothesized that differences in the specification of gametophytic cells could predetermine this. Nevertheless, the regulatory control is not always strict, as unfrequently fertilization of an unreduced egg cell in apomictic gametophytes takes place and leads to ploidy duplication in the BIII hybrids formed (Hojsgaard and Hörandl, 2019).

In contrast to *A. thaliana*, in *Boechera FAS4* is represented by 3 genes. It is tempting to speculate that they might be involved in establishing preconditions to allow double fertilization in sexual plants versus single fertilization in pseudogamous apomicts. In this line, whole genome duplications and small duplications restricted to smaller genomic regions are drivers of evolution, as a relieve of selection pressure lowers purifying selection and allows accumulation of mutations for new gene copies (Carretero-Paulet and Fares, 2012). This can either result in loss of function or allow functional diversification leading to sub-or neo-functionalization. In addition, regulatory elements in promoter regions or similar regulatory mechanisms can lead to alterations in expression thereby promoting changes in regulation and subsequently function (Singh and Hannenhalli, 2008). Strikingly, in case of *FAS4* in *Boechera*, not the orthologue *0032*, but the derived paralogue *0021* shows higher expression in mature ovules as compared with earlier stages of development consistent with the expectations for *0032*, and higher expression in sexual as compared to apomictic ovules. Phylogenetic analysis revealed that both genes form distinct clusters. Interestingly, alleles from facultative apomictic or apomictic accessions largely form a different cluster than alleles from sexual accessions for *0021*, unlike for *0032*. It can therefore be hypothesized that the distinct copies adopted different roles in shaping of reproductive development and aspects of gametogenesis. Nevertheless, alleles of two apomictic accessions (ES517 and C011010) do not cluster together with alleles of other (facultative) apomictic accessions. Possibly, the regulatory control they are involved in is not strictly the same as in other apomictic accessions. This might relate to low frequencies of sexuality observed, at least for ES517 (Mau et al., 2015). Future studies on a possible correlation of frequencies of BIII hybrids formed per accession and functional investigations are required to gain insights into the roles of the distinct *FAS4* genes.

In addition, tight molecular control mechanisms are required to allow autonomous activation of the egg cell synchronized with initiation of endosperm development. Our comprehensive transcriptional analysis identified a number of candidate genes potentially involved in this process. In good agreement with differences of the epigenetic makeup of sexual and apomictic gametes, we identified the homologues of *CHR34* and *JMJ27* as higher expressed in sexual as compared to apomictic ovules. Interestingly, in *A. thaliana* JMJ27 is essential for male meiosis (Cheng et al., 2022), but roles during later stages of reproductive development have not been reported. In sexual species, egg cells are characterized by a repressive state of chromatin based on histone marks such as H3K9, whereas parthenogenesis might rely on a more permissive state to overcome this developmental block. As JMJ27 removes repressive methylation marks (Dutta et al., 2017), it can be hypothesized that deposition of translationally silenced mRNAs of this gene might allow its rapid activation upon fertilization resulting in fast removal of this repressive mark. However, to date, no studies supporting this or potential roles of CHR34 in modulation of chromatin structure have been reported. Interestingly, the set of genes expressed at higher levels in the ovules of apomicts than of sexual *Boechera* included *KRP7*. KRP7 acts redundantly with other KIP-proteins to control cell cycle progression and aspects of reproductive development by repression of the cell cycle regulator Retinoblastoma homologue *RBR1* (Zhao et al., 2017, Verkest et al., 2005). Therefore, the higher abundance in samples from apomictic as compared to sexual ovules might be aiding the activation of cell cycle activity as a consequence of gradual downregulation of KRP activity (Sizani et al., 2019). Interestingly, also homologues of *ESP3* are differentially regulated at distinct stages of reproductive development and between sexual and apomictic plants. The phylogenetic close relation to *MEE29* and an additional so far uncharacterized gene, as well as the identification of Bostr.*7867s0308* as DEG which has no homologue in *A. thaliana* suggests that a complex evolutionary history of the genes might be relevant for shaping the plasticity of reproductive systems. In addition, a homologue of *RCK* was higher abundant in apomictic as compared to sexual ovules. As RCK is a meiotic gene involved in crossover formation (Chen et al., 2005), the functional relevance of differential abundance in mature gametophytes is questionable. Previously, several meiotic genes have been identified to be present in the AIC of the triploid apomict *B. gunnisoniana* (Schmidt et al., 2014), consistent with this cell to initiate but alter meiosis. As meiotic recombination is skipped in apomicts, the identification of *RCK* in mature ovules might either represent stored mRNA transcript that has not been translated and was not yet subjected to degradation, or the *Boechera* homologue might serve roles in mitotic recombination potentially relevant for prevention of mutation accumulation in apomicts. In summary, our study strengthened the evidence that diverse RNA helicases are crucial for plant germline development and aspects of their evolution can shape the plasticity of plant reproduction.

## Methods

### Primers used

All primers used in this study are listed in Table S11.

### Plant material

Seeds of *Boechera* Á.Löve & D.Löve were obtained as previously described (Kiefer et al., 2020). Cultivation and *Boechera* plants and ploidy analysis of apomictic plants was done as described previously (Zühl et al., 2019, Kiefer et al., 2020). For functional analysis *Arabidopsis thaliana* (L.) Heynh. was used. Ecotype Col-0 used as wild-type and for generating CRISPR/Cas9 lines. Seeds of the p*WOX2::CENH3-GFP* marker line (De Storme et al., 2016) and the p*AKV::H2B-YFP* marker line (Rotman et al., 2005) were kindly provided by Danny Geelen (Ghent University, Ghent, Belgium) and Weicai Yang (Beijing University, Beijing, China), respectively. The T-DNA insertion line SAIL_324_E12 in *At1G33390* (referred to as *fas4-1*) was ordered from “The Nottingham Arabidopsis Stock Centre” (NASC, arabidopsis.info). The insertion site was confirmed to be in the 4^th^ intron (Supporting Information). Genotyping was performed with primers LB, LP and RP. Selection of seedlings was done on murashige-skoog (MS) plates supplied with 10 µg/ml phosphinothricin. Seedlings were germinated and grown on MS plates for about 2 weeks before transfer to soil (ED73, Universalerde, Germany). Plants were grown in a growth chamber at 16h light / 8h darkness at 21°C and 18°C and treated with nematodes against black flies, respectively. Plants harbouring the *AtFAS4*_genomic_-*PmTurquoise* or *BsFAS4orth* _genomic_-*mVenus* construct were cultivated in an incubator at 22°C at 14h light/ 10 h darkness or in a greenhouse.

### Generation of CRISPR/Cas9 line

To generate *fas4-2, fas4-3*, and *fas4-4*, small guiding RNAs (sgRNAs) were designed using the online tool chopchop (https://chopchop.cbu.uib.no/). The oligonucleotides 5’-ATTGGAAAGTAGTAGCAAGCTTG-3’ and 5’-AAACCAAGCTTGCTACTACTTTC-3’ were ligated and introduced into the vector pHEE401E (Wang et al., 2015). Generation of lines was further done following established protocols with minor modifications (Supporting Information) (Kiefer et al., 2020).

### Generation of marker lines for orthologue expression

As template for amplification of the constructs, genomic DNA was extracted from leaves of *A. thaliana* Col-0 plants and *B. stricta* LTM with Invisorb Spin Plant Mini Kit (Invitek Molecular GmbH, Berlin, Germany) following manufacturer instructions. Cloning was done with Polymerase Incomplete Primer Extension (PIPE) cloning (Klock and Lesley, 2009). Constructs pEarlyGate201-AtFAS4-PmTurquoise and pEarlyGate201-BsFAS4ortho-mVenus, the expression constructs of *AtFAS4*_genomic_-*PmTurquoise* and *BsFAS4orth*_genomic_-*mVenus*, respectively, harbour genomic regions from the *A. thaliana* and *B. stricta* LTM orthologues in translational fusion to the genes encoding for PmTurquoise and mVenus, respectively (Supporting Information). The final constructs were transformed into *Agrobacterium tumefaciens* strain GV3101 before transforming *fas4-1/FAS4* plants by floral dip (Clough and Bent, 1998). Transformants were selected on MS plates containing phosphinothricin and tested for presence of the construct with primers P23 and P24, for the presence or absence of the insertion in *FAS4* with the LB and RP primer combination, and for testing for a wild-type allele of the *FAS4* locus the LP was used together with P25. For all genotyping amplification reactions Phire Hot Start II DNA Polymerase (Thermo Fisher Scientific) was applied.

### Isolation of reproductive tissues for RNA-Seq and library preparation

Isolation of nucellus tissues from *Boechera* with LAM, subsequent RNA isolation, quality control, and amplification was performed as previously described (Supporting Information) (Zühl et al., 2019). Per sample, between 99 and 223 sections of nucellus tissue were pooled (Table S1). Isolation of mature ovules and subsequent RNA isolation followed protocols previously described for *A. thaliana* (Stein et al., 2021) (Supporting Information). Library preparation from samples isolated by LAM was done with NEB Ultra II FS gDNA library preparation kit. In addition to the samples generated in this study, libraries were prepared from stored cDNAs of samples generated previously (Zühl et al., 2019). Libraries of ovule samples were prepared with the NEBNext Ultra II Directional RNA Library Prep Kit for Illumina (New England Biolabs) using NEBNext® Multiplex Oligos for Illumina (Index Primers Set 1 and Index Primers Set 2) for sample indexing. Sequencing was performed on an Illumina NextSeq 500 instrument (Illumina, San Diego, USA) by the Deep Sequencing Core Facility of Heidelberg University using the 75 bp PE protocol. The quality of all libraries was confirmed using Qubit (Thermo Fisher Scientific) and Bioanalyzer High Sensitivity DNA assays.

### Data analysis

#### Quality control and trimming of raw reads

Quality control and trimming of raw reads was done as previously described (Zühl et al., 2019). STAR version 2.5.3a_modified was used for read mapping using the *B. stricta* LTM genome assembly and annotation (Lee et al., 2017, Dobin et al., 2013). The number of fragments of correctly paired reads, of which minimum 70% of read length mapped to an exon, was determined using featureCounts in Rsubread package (version 1.20.6), as described previously (Liao et al., 2014, Schmid, 2017, Zühl et al., 2019).

#### Differential expression analysis and gene ontology analyses

Differential expression analysis was done with the Bioconductor package EdgeR implemented in R (Robinson et al., 2009). For statistical analysis all samples from accessions of the same reproductive mode and of the same developmental stage (i.e. nucellus tissues harbouring the MMC or AIC, nucellus tissues harbouring the FMS, ovule tissues harbouring mature gametophytes) were treated as biological replicates. For ANOVA-like analysis of transcriptional changes at different stages of development, first all samples from sexual or apomictic samples were selected. EdgeR was used for filtering of low expressed genes and estimation of dispersion, before applying a generalized linear model and a quasi-likelihood F-test (Robinson et al., 2009). The design matrix used for the developmental stages as factor levels and included an intercept. A pairwise comparison was done on all samples from nucellus tissues harbouring MMCs collected from sexual accessions versus all ovule samples from sexual accessions. In addition, a pairwise comparison was performed to identify DEGs between ovules from sexual accessions and ovules from apomictic accessions. The pairwise comparisons were done using the classic edgeR pipeline (Robinson et al., 2009). DEGs were defined to show an FDR < 0.05 after Benjamini-Hochberg adjustment.

#### Gene ontology enrichment analysis

For gene ontology enrichment analyses we first identified the *A. thaliana* homologues of the *Boechera* genes based on the annotation in the reference genome (Lee et al., 2017). To test for enrichment of molecular functions and biological processes the Bioconductor package topGO was used (Alexa A, 2019). As customized gene to GO annotations (gene2GO) we downloaded the ATH_GO_GOSLIM annotations from TAIR (arabidopisis.org). To test for overrepresented GO terms Fisher’s exact test was used combined with the function “weight”.

#### Generation of Venn diagram and Heatmaps

The heatmap.2 function implemented in the Bioconductor package gplots (Warnes et al., 2005) was used to generate the heatmap using agglomerative clustering (complete linkage) and euclidean distance. Heatmaps were based on log2-transformed read counts after TMM-normalization done with NOISeq implemented in R (Tarazona et al., 2015). The Venn diagram was generated using the online tool InteractiVenn (Heberle et al., 2015).

#### Gene evolution of ESP3 and MEE29 homologues

Blast search of the protein sequence of *At*ESP3 using standard settings in TAIR (www.arabidopsis.org) was applied to identify high similarities between *At*ESP3 and *At*MEE29. Blast search on Phytozome v13 (https://phytozome.jgi.doe.gov) was used for detecting sequences orthologous or paralogous to *ESP3* and *MEE29* in *Euclidium syriacum* (Brassicales Map Alignment Project, DOE-JGI, http://bmap.jgi.doe.gov/), *Arabis alpina* (Willing et al., 2015), *Arabidopsis lyrata* (Hu et al., 2011, Rawat et al., 2015), *Eutrema salsugineum* (Yang et al., 2013) and *Arabidopsis thaliana* (Cheng et al., 2017), as well as *Boechera stricta v1*.*2* (DOE-JGI, http://phytozome.jgi.doe.gov/). Neighbouring genes were inspected in the gbrowser and checked for their closest orthologue in *A. thaliana*. It was not possible to determine positional orthologues in all taxa as in some cases duplication events were found to be lineage specific. For *Arabis alpina*, the neighbouring genes were checked in the genome browser available on arabis-alpina.org. The orthologue list deposited there was used to identify the *A. thaliana* orthologues of neighbouring genes (Willing et al., 2015). For phylogenetic reconstruction all CDS were extracted and aligned by mafft (Katoh et al., 2002). The generated alignment was used as input for raxml-ng (Kozlov et al., 2019) where GTR+I+G was set as model and 500 bootstrap replicates were run. For determining possible conserved blocks upstream and downstream as well as in the CDS of *ESP3* and *MEE3* orthologues and related sequences, sequence stretches reaching from the end of the upstream gene until the beginning of the downstream gene were extracted using bedtools getfasta (Quinlan and Hall, 2010). An mVISTA analysis(Frazer et al., 2004) using LAGAN (Brudno et al., 2003) for alignment was run on the extracted sequences. The resulting plots were further edited (colour, font size, labeling) in Adobe Illustrator for better comprehension.

#### Gene evolution of FAS4 orthologues and paralogues and generation and analysis of phased FAS4 alleles

Inspection of the *B. stricta LTM* draft genome (Lee et al., 2017) revealed it contains three sequences showing high homology to *A. thaliana FAS4*. In order to obtain a better understanding of *FAS4* evolution, several published Brassicaceae draft and reference genomes (*Euclidium syriacum* (Brassicales Map Alignment Project, DOE-JGI, http://bmap.jgi.doe.gov/), *Arabidopsis lyrata* (Hu et al., 2011, Rawat et al., 2015), *Eutrema salsugineum* (Yang et al., 2013), *Arabidopsis thaliana* (Cheng et al., 2017) as well as *Boechera stricta v1*.*2* (DOE-JGI, http://phytozome.jgi.doe.gov/) were queried by blast search on Phytozome v13 (https://phytozome.jgi.doe.gov) using the genic sequences of *AtFAS4* (*At1g33390*) as well as the upstream and downstream genes *At1g33360* and *At1g33400*. If the three genes were not syntenic, the neighboring genes of the blast hit for *FAS4* were queried against the *A. thaliana* genome (www.arabidopsis.org). For *Arabis alpina, FAS4* as well as neighbouring genes were looked up in a 1:1 orthologue list by their *Arabidopsis* gene number (Willing et al., 2015). Sequences spanning the region between *FAS4* orthologues or paralogues and their respective upstream and downstream genes as well as mVISTA analysis was carried out as described above for *ESP3* and *MEE29*.

For generating phased alleles of the *FAS4* positional ortholog as well as of the paralogues sequences on scaffold 17016 the reads from the previously published gene capture experiment were re-analysed. Reads were trimmed using Trimmomatic 0.22 (Bolger et al., 2014) and mapped to the three loci identified as possible *FAS4* homologues including also all upstream and downstream sequence until the next gene using bwa mem (Li, 2013, Li and Durbin, 2009). Mapping quality was enhanced using samtools (Li et al., 2009) and picardtools (http://broadinstitute.github.io/picard) was used for removing duplicates. Variant calling was performed using gatk4 HaplotypeCaller. The resulting variants along with the quality enhanced output from read mapping were used as input for WhatsHap (Patterson et al., 2015) which was used for haplotype phasing. The output of WhatsHap was inspected for sequences which could be phased in one block and haplotype sequences were reconstructed using bcftools consensus. Sequences which were phased in two blocks were inspected for length of the phased blocks. If all CDS was covered, these sequences were also included in the analysis, excluding the unphased part. In a last step low coverage regions or regions with low mapping quality were identified by gatk38 CallableLoci (McKenna et al., 2010) and masked by using bedtools maskfasta (Quinlan and Hall, 2010).

As phased alleles could only be obtained for the positional ortholog as well as for gene 0021 on scaffold 17016, further analyses only included these two loci and gene *0022* on scaffold 17016 of the reference accession LTM (from (Lee et al., 2017)). Sequences for each locus were aligned to the sequences used as reference in mapping by using MAFFT (Katoh et al., 2002) and exons were extracted manually by examining and cutting the alignment when visualized in aliview (Larsson, 2014). The resulting continuous coding sequences were translated using transeq (https://www.ebi.ac.uk/Tools/st/emboss_transeq) and realigned using MAFFT. Frameshifts and stop codons were identified.

A combined set of all phased coding sequences for *0032* and *0021* as well as the reference sequences for *0032, 0021* and *0022* and the CDS of *AtFAS4* was also aligned using MAFFT. jModeltest (Posada, 2008) was used for determining the best molecular evolutionary model (HKY + G) and raxml-ng (Kozlov et al., 2019) was used for phylogenetic reconstruction including 500 bootstrap replicates.

#### Analysis of pollen viability and developmental defects

To analyse pollen viability we followed a modified protocol for Alexander stain (Ross et al., 2010). 4’,6-diamidino-2-phenylindole (DAPI) staining was done on mature pollen shortly after anthesis as previously described using a final concentration of 1µg/ml DAPI (Backues et al., 2010).

#### Clearing and microscopy

To investigate female gametophyte development in the *A. thaliana* lines carrying mutant alleles of *FAS4* as compared to the wild-type, ovule clearing and microscopy were performed as described previously (Stein et al., 2021) (Supporting Information), unlike that for *AtFAS4*_genomic_-*PmTurquoise* or *BsFAS4orth* _genomic_-*mVenus* lines imaging were acquired sequentially with a Zeiss LSM710 microscope.

## Acknowledgement

We are grateful to Marcus A. Koch (Centre for Organismal Studies (COS), Heidelberg, Germany) for sharing the working time of his technical assistant. We thank the groups at COS (Heidelberg University), Joachim P. Spatz (Max Planck Institute for Medical Research, Germany), and Philipp Schlüter (University of Hohenheim, Germany) for generous access to equipment and facilities, the Technology Platform “Cellular Analytics” of the Stuttgart Research Center Systems biology for their support, and the people from Phytotechnikum Hohenheim for access to facilities and help with plant care. We thank George Coupland (Max Planck Institute for Plant Breeding Research, Cologne, Germany) for providing vectors Tmp-pSTB205 and pEarlyGate301, Danny Geelen (Ghent University, Ghent, Belgium) and Weicai Yang (Beijing University, Beijing, China) for providing marker lines, Timothy Sharbel (Global Institute for Food Security, Saskatoon, Canada), Thomas Mitchell-Olds (Duke University, USA), Ueli Grossniklaus (University of Zürich, Switzerland) and John Carman (Utah State University, USA) for providing *Boechera* seeds and we are grateful to the staff of the Botanical Garden (Heidelberg, Germany) for seed propagation. We also acknowledge DFG for funding (SCHM2448/2-1 and SCHM2448/2-2) to AS.

## Data availability

RNA-Seq data are deposited on the NCBI Sequence archive database under number PRJNA863097.

**Figure S1**: Heatmap of expression of 71 *Boechera* homologues of RNA helicases. The helicases were identified as DEGs in an ANOVA-like analysis comparing different stages of reproductive development in sexual accessions. The heatmap is based on log2-scale normalized read counts of all samples from sexual accessions of nucellus tissues harbouring the MMC, and of mature ovules. Hierarchical clustering of genes/samples was based on euclidian distance and hierarchical agglomerative clustering. Colors are scaled per row. Red denotes high and black low expression.

**Figure S2**: Heatmap of expression of 917 genes differentially expressed in ovules from sexual as compared to apomictic accessions. The heatmap is based on log2-scale normalized read counts. Hierarchical clustering of genes/samples was based on euclidian distance and hierarchical agglomerative clustering. Colors are scaled per row. Red denotes high and black low expression.

**Figure S3: Expression of *Arabidopsis* gametophyte enriched genes and RNA helicases**. Shown are such genes identified as DEGs in sexual as compared to apomictic ovules harbouring mature gametophytes. Gene identifiers are given for *B. stricta* LTM, in addition to gene names or identifiers of homologues in *A. thaliana*. The heatmap is based on log2-scale transformed TMM-normalized read counts. Hierarchical clustering of genes and samples was applied based on euclidean distance and hierarchical agglomerative clustering. Colors are scaled by row with red denoting high and black denoting low expression.

**Figure S4**: Evolutionary analysis of *ESP3, MEE29* and homologues identified *Bostr*.*3359s0111* as the positional orthologue of *AtESP3*, and *Bostr*.*30275s126* as orthologue of *At4G16680*, whereas no *A. thaliana* orthologue was found for *Bostr*.*7867s0308*.

**Figure S5**: Phylogenetic analysis of homologues of *ESP3, MEE29* and related genes by maximum likelihood analysis using RAxML (Stamatakis, 2014).

**Figure S6**: Vista plots indicating similarities of orthologues from *ESP3* (A), *MEE29* (B), *At4G16680* (C), and *Bostr*.*7867s0308* in genomes of six selected Brassicaceae species using sequences from *A. alpina* (A,C,D) and *E. salsugineum* as base.

**Figure S7**: Phylogenetic analysis of homologues of *FAS4* by maximum likelihood analysis using RAxML (Stamatakis, 2014).

**Figure S8**: Localization of *At*FAS4 and *Bs*FAS4 orthologues in reproductive tissues using confocal laser scanning confocal microscopy. (A-D) Signals from fluorescence of PmTurquoise in plants carrying the *AtFAS4*_genomic_-*PmTurquoise* construct in developing ovule harbouring a developing gametophyte (A), mature gametophyte (B,C), and in anther tissues (D). (E,F) Signal from fluorescence of mVenus in anther tissues harbouring the and *BsFAS4orth* _genomic_-*mVenus*. 4nuc, 4 nucleate gametophyte; gam, mature gametophyte; syn, synergids; the arrow depicts a nucleus in anther tissues; * indicate pollen; scales = 20 µm.

**Figure S9**: Clearing of *A. thaliana* ovules at 2 days after emasculation observed by DIC microscopy. (A,B) Wild-type ovules with mature gametophyte (A), and gametophyte with unfused polar nuclei (B). (C-J) Ovules from *fas4-1/FAS4* harbouring a wild-type mature gametophyte (C), an developmentally delayed or arrested 2-nucleate gametophyte (D), aberrant 2-nucleate gametophytes (E,F), a 4-nucleate gametophyte (I), and gametophytes with aberrant numbers of gametophytic nuclei (G,H,J). cc, central cell; egg, egg cell; syn, synergid cells; nuc, nucleate. Scale bars are 50 µm.

**Figure S10**: Developmental defects during female gametogenesis in *fas4-1/FAS4*. (A-E) Laser scanning confocal microscopy of developing ovules of plants harbouring the p*AKV::H2B-YFP* (Rotman et al., 2005) construct. Wild-type like FMS (A), 2-nucleate gametophyte (2nuc) at early (B) and later stage of development (C), and 4-nucleate gametophyte (4nuc) (D). (E) Mutant gametophyte showing an aberrant number of 3 gametophytic nuclei. (F-J) Epifluoresence microscopy on developing ovule carrying the p*WOX2::CENH3-GFP* marker (De Storme et al., 2016) for visualization of chromosomes in gametophytic nuclei. (F,G) Wild-type like gametophyte with 5 dots demarking the chromosomes in the haploid gametophytic lineage per nucleus in the FMS (F) and the 2-nucleate (2nuc) gametophyte (G). (H-J) Mutant gametophytes showing aberrations in the division plane/localization of gametophytic nuclei after the first mitotic division (H), alterations in migration (I), or separation of gametophytic nuclei (J). Scale bars = 20 µm.

**Figure S11**: Seedlings of *fas4-1/FAS4* (B,D) show significantly increased resistance to streptomycin as compared to wild-type (A,C). (A-D) 14 day old seedlings germinated and grown on murashige-skoog medium without (A,B), or supplied with 30 µg ml^-1^ streptomycin (C,D). (E) Whereas no significant difference was observed concerning approximate rosette diameter in mm for 14 days old seedlings not supplied with streptomycin (gray boxes), on medium supplied with streptomycin rosette diameter were significantly larger for lines carrying a mutant allele of FAS4 as compared to wild-type (white boxes, *** indicates p < 0.005 as tested with two-sided students t-test). Scales = 1 cm.

**Table S1**: Summary of samples, input material, and mapping statistics. Given are sample names, sequencing technology with respect to single end (SE) or paired end (PE) sequencing, numbers of sections isolated by laser assisted microdissection (if applicable), the developmental stage of the samples, numbers of total raw reads and numbers of (paired) read counts as analysed by feature counts after quality control, trimming, and mapping. * indicates that libraries were constructed from cDNA generated in (Zühl et al., 2019).

**Table S2**: Analysis of differential gene expression at different developmental stages in sexual (A) and apomictic (B) *Boechera*. Differentially expressed genes were identified in an ANOVA-like analysis using quasi-likelihood F-test implemented in edgeR using a generalized linear model (Robinson et al., 2009). Group sex2 or apo2 thereby tests for differences between nucellus tissues harbouring MMC/AIC stage, and sex3 or apo3 tests for differences between MMC/AIC stage and mature ovules. Given are log fold changes (FC) per group, F-values, and p-values. Genes identified at FDR≤0.05 were considered significant.

**Table S3**: Gene ontology analysis to identify enriched molecular functions based on 9‘136 genes identified as differentially regulated at different stages of development in apomictic *Boechera* (p < 0.01).

**Table S4**: Analysis of differential expression between all samples from sexual accessions at MMC stage and mature ovules (FDR≤0.05). Classical pairwise comparisons were applied using edgeR (Robinson et al., 2009).

**Table S5**: Gene ontology analysis. Enrichment of biological processes was identified based on 4‘219 gene upregulated in nucellus tissues harbouring MMCs (A), and 3‘907 genes upregulated in mature ovule tissues (B) (p < 0.01). Upregulated genes were identified in pairwise comparison of differential expression.

**Table S6**: RNA helicases identified as differentially expressed at different developmental stages in sexual *Boechera*. Raw read counts in any sample and annotations are given for the 71 homologues of RNA helicases identified as DEGs.

**Table S7**: Genes differentially expressed between mature ovules of all sexual as compared to all apomictic *Boechera* accessions. Significantly differential expression was identified in a pairwise comparison using edgeR (FDR≤0.05) (Robinson et al., 2009). Non-normalized read counts are given.

**Table S8:** Microsynteny and genomic regions of FAS4 homologues, ESP3 and MEE29 and related genes. Directionalities of the genes in the genomes are indicated by arrows.

**Table S9:**
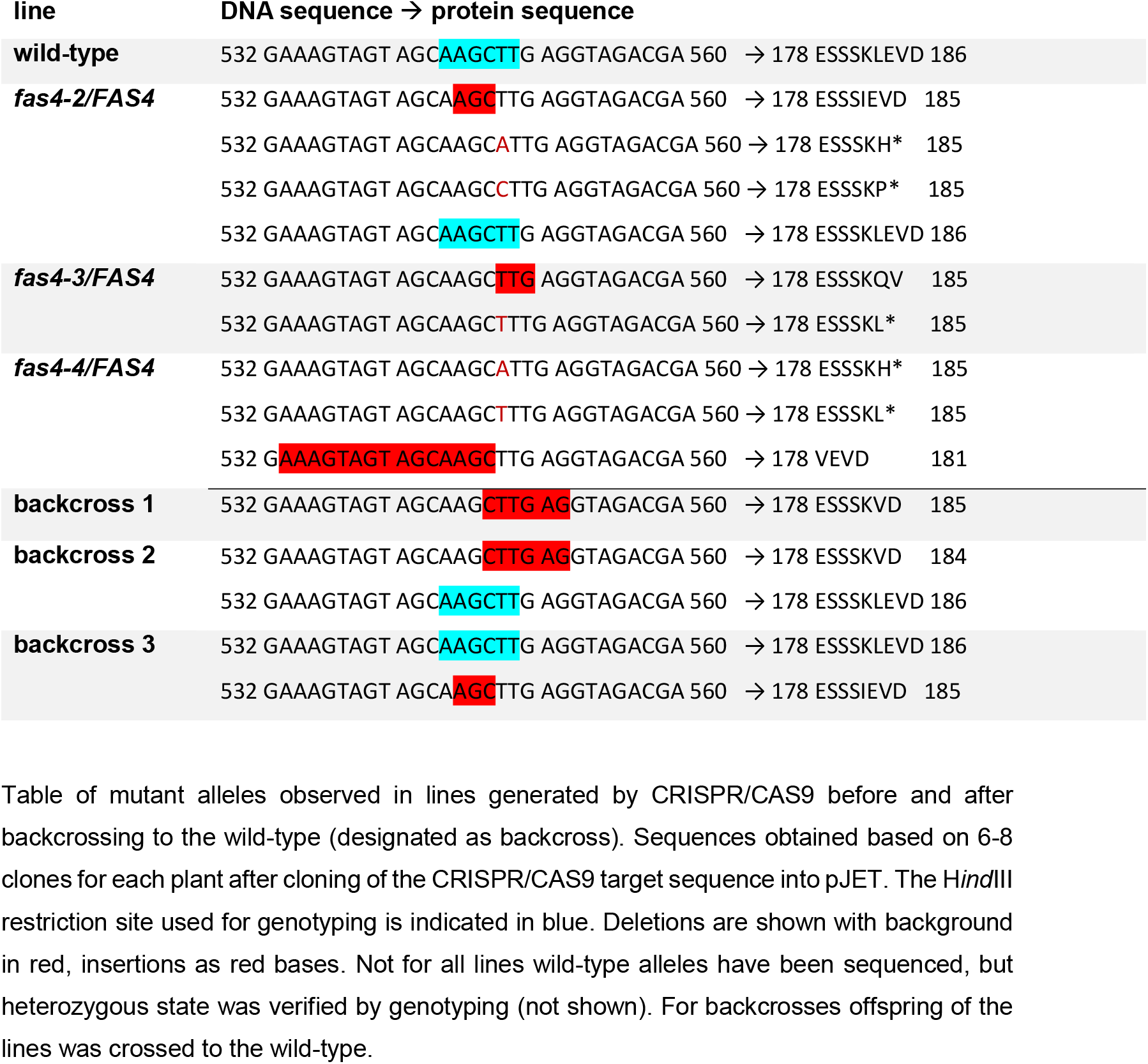
Mutant alleles identified in CRISPR/CAS9 lines.

**Table S10:** Phenotypic defects observed in *fas4-1/FAS4* harbouring the p*AKV::H2B-YFP* marker for labelling of gametophytic nuclei.

| Developmental stage of most advanced gametophytes (wild-type like) | FMS | 2-nucleate gametophyte (wild-type like) | 4-nucleate gametophyte (wild-type like) | mature gametophyte (wild-type like) | 2-nucleate gametophyte (odd) | aberrant numbers or positions of gametophytic nuclei | n total |
| --- | --- | --- | --- | --- | --- | --- | --- |
| **FMS-2-nucleate** | 48.5 | 46.2 | 0 | 0 | 5.3 | 0 | 132 |
| **2-/4-nucleate** | 28.3 | 32.9 | 7.9 | 0 | 18.4 | 12.5 | 152 |
| **mature** | 8.5 | 13.6 | 18.6 | 16.9 | 8.5 | 33.9 | 59 |

Data were collected and summarized from 5 plants and demonstrates that developmental defects were mainly observed after the first mitotic division.

**Table S11:**
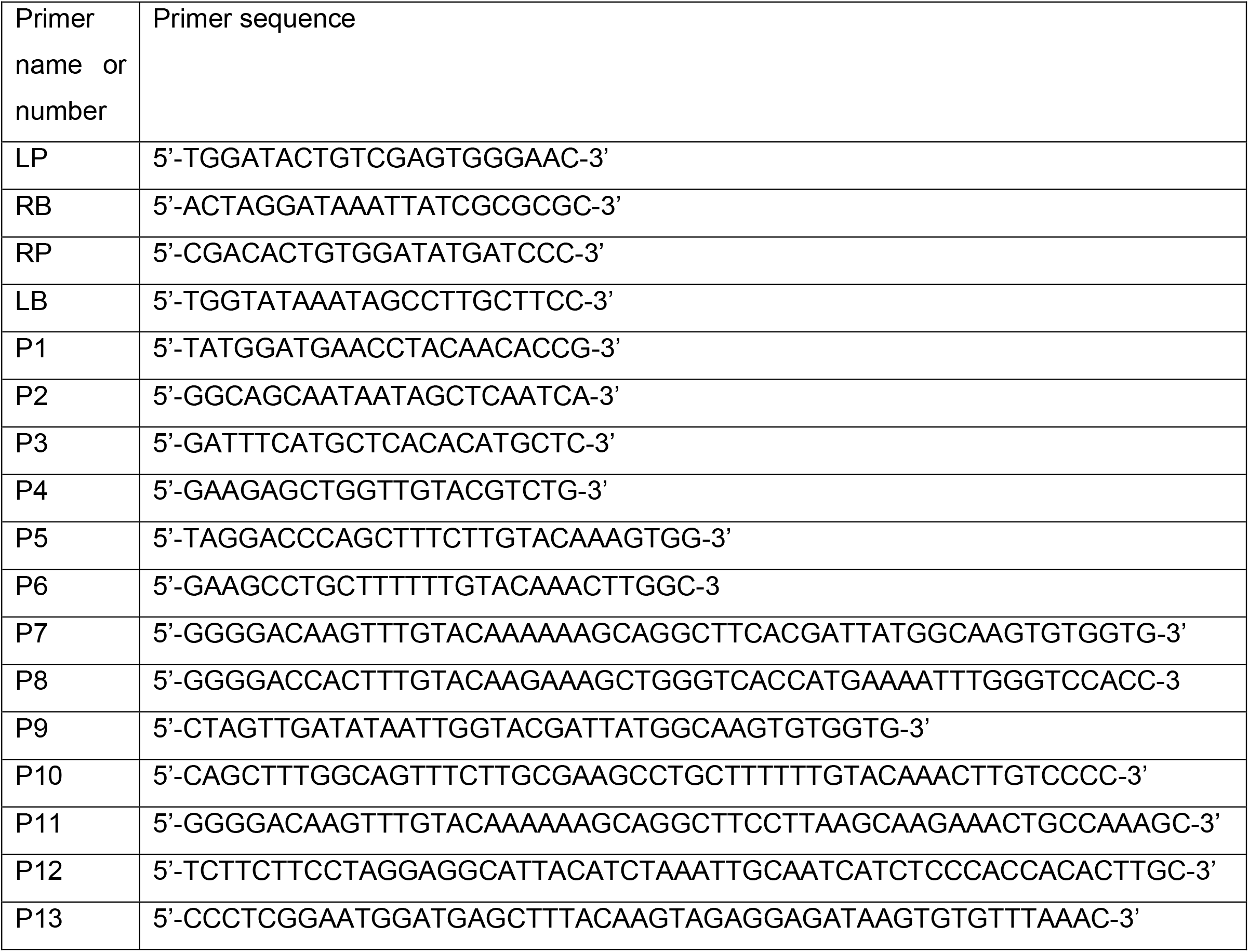
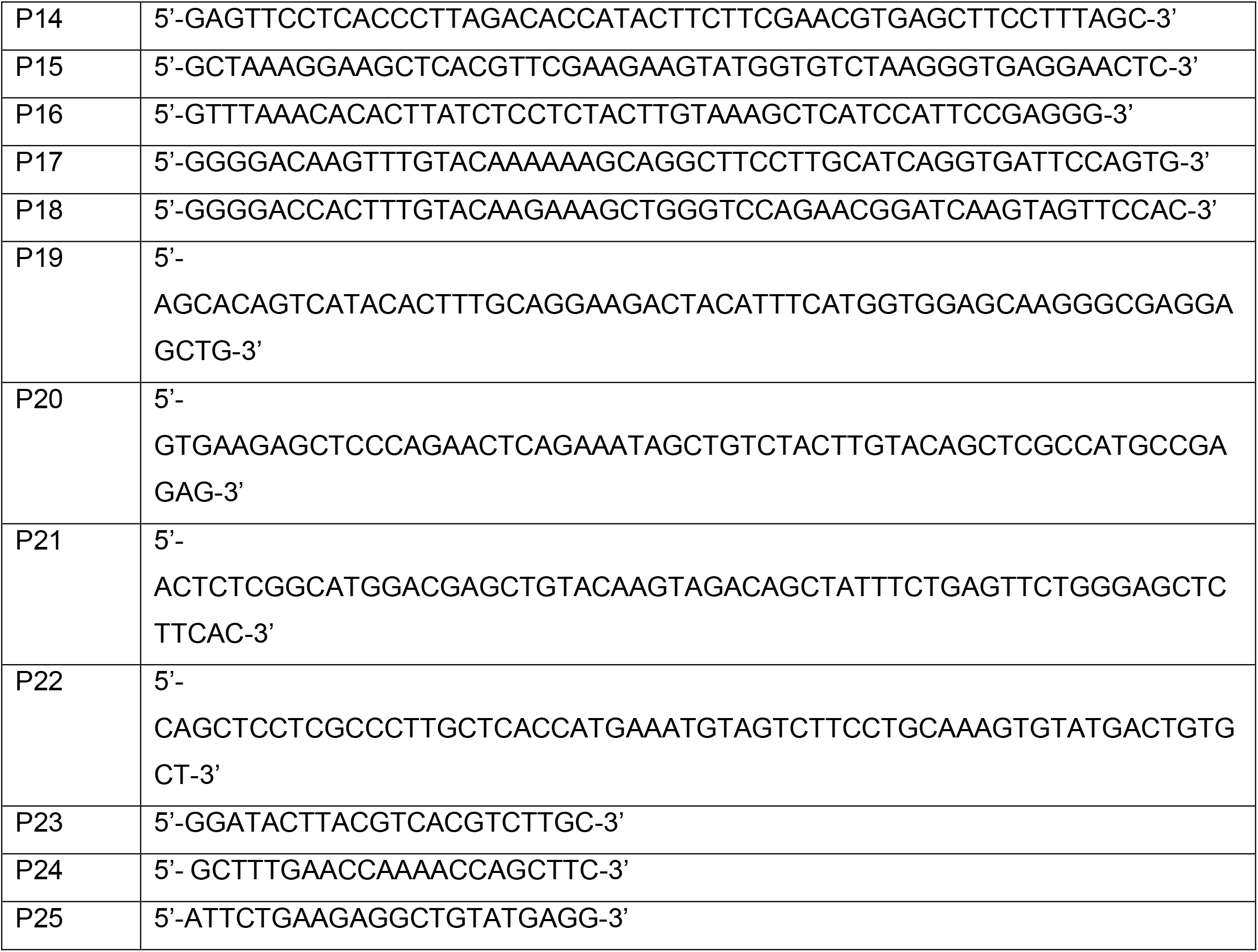
Table of primers used.

## References

Alexa A R. J. 2019. topGO: Enrichment Analysis for Gene Ontology. R package version, 2.38.1.

Aliyu, O. M., Schranz, M. E. & Sharbel, T. F. 2010. Quantitative variation for apomictic reproduction in the genus Boechera (Brassicaceae). Am J Bot, 97, 1719–31.

Backues, S. K., Korasick, D. A., Heese, A. & Bednarek, S. Y. 2010. The Arabidopsis dynamin-related protein2 family is essential for gametophyte development. Plant Cell, 22, 3218–31.

Bansal, P., Madlung, J., Schaaf, K., Macek, B. & Bono, F. 2020. An interaction network of RNA-binding proteins involved in Drosophila oogenesis. Mol Cell Proteomics, 19, 1485–1502.

Barak, S., Singh Yadav, N. & Khan, A. 2014. DEAD-box RNA helicases and epigenetic control of abiotic stress-responsive gene expression. Plant Signal Behav, 9, e977729–e977729.

Barcaccia, G. & Albertini, E. 2013. Apomixis in plant reproduction: a novel perspective on an old dilemma. Plant Reprod, 26, 159–79.

Bolger, A. M., Lohse, M. & Usadel, B. 2014. Trimmomatic: a flexible trimmer for Illumina sequence data. Bioinformatics, 30, 2114–20.

Brudno, M., Do, C. B., Cooper, G. M., Kim, M. F., Davydov, E., Green, E. D., Sidow, A. & Batzoglou, S. 2003. LAGAN and Multi-LAGAN: efficient tools for large-scale multiple alignment of genomic DNA. Genome Res, 13, 721–31.

Carretero-Paulet, L. & Fares, M. A. 2012. Evolutionary Dynamics and Functional Specialization of Plant Paralogs Formed by Whole and Small-Scale Genome Duplications. Mol Biol Evol, 29, 3541–3551.

Chen, C., Zhang, W., Timofejeva, L., Gerardin, Y. & Ma, H. 2005. The Arabidopsis ROCK-N-ROLLERS gene encodes a homolog of the yeast ATP-dependent DNA helicase MER3 and is required for normal meiotic crossover formation. Plant J, 43, 321–34.

Cheng, C.-Y., Krishnakumar, V., Chan, A. P., Thibaud-Nissen, F., Schobel, S. & Town, C. D. 2017. Araport11: a complete reannotation of the Arabidopsis thaliana reference genome. Plant J, 89, 789–804.

Cheng, J., Xu, L., Bergér, V., Bruckmann, A., Yang, C., Schubert, V., Grasser, K. D., Schnittger, A., Zheng, B. & Jiang, H. 2022. H3K9 demethylases IBM1 and JMJ27 are required for male meiosis in Arabidopsis thaliana. New Phytol, 253, 2252–2269.

Clough, S. J. & Bent, A. F. 1998. Floral dip: a simplified method for Agrobacterium-mediated transformation of Arabidopsis thaliana. Plant J, 16, 735–43.

Conner, J. A. & Ozias-Akins, P. 2017. Apomixis: Engineering the Ability to Harness Hybrid Vigor in Crop Plants. Methods Mol Biol, 1669, 17–34.

De Storme, N., Keçeli, B. N., Zamariola, L., Angenon, G. & Geelen, D. 2016. CENH3-GFP: a visual marker for gametophytic and somatic ploidy determination in Arabidopsis thaliana. BMC Plant Biol, 16, 1.

Dobin, A., Davis, C. A., Schlesinger, F., Drenkow, J., Zaleski, C., Jha, S., Batut, P., Chaisson, M. & Gingeras, T. R. 2013. STAR: ultrafast universal RNA-seq aligner. Bioinformatics (Oxford, England), 29, 15–21.

Dutta, A., Choudhary, P., Caruana, J. & Raina, R. 2017. JMJ27, an Arabidopsis H3K9 histone demethylase, modulates defense against Pseudomonas syringae and flowering time. Plant J, 91, 1015–1028.

Ebel, C., Mariconti, L. & Gruissem, W. 2004. Plant retinoblastoma homologues control nuclear proliferation in the female gametophyte. Nature, 429, 776–780.

Frazer, K. A., Pachter, L., Poliakov, A., Rubin, E. M. & Dubchak, I. 2004. VISTA: computational tools for comparative genomics. Nucleic Acids Res, 32, W273–9.

Friedman, W. E. & Williams, J. H. 2003. Modularity of the angiosperm female gametphyte and its bearing on the early evolution of endosperm in flowering plants. Evolution, 57, 216–230.

Grimanelli, D. 2012. Epigenetic regulation of reproductive development and the emergence of apomixis in angiosperms. Curr Opin Plant Biol, 15, 57–62.

Heberle, H., Meirelles, G. V., Da Silva, F. R., Telles, G. P. & Minghim, R. 2015. InteractiVenn: a web-based tool for the analysis of sets through Venn diagrams. BMC Bioinformatics, 16, 169.

Herr, A. J., Molnàr, A., Jones, A. & Baulcombe, D. C. 2006. Defective RNA processing enhances RNA silencing and influences flowering of Arabidopsis. Proc Nat Acad Sci U S A, 103, 14994–15001.

Hojsgaard, D. & Hörandl, E. 2019. The Rise of Apomixis in Natural Plant Populations. Front Plant Sci, 10, 358.

Honys, D., RĔnák, D., Feciková, J., Jedelský, P. L., Nebesárová, J., Dobrev, P. & Capková, V. 2009. Cytoskeleton-associated large RNP complexes in tobacco male gametophyte (EPPs) are associated with ribosomes and are involved in protein synthesis, processing, and localization. J Proteome Res, 8, 2015–31.

Hörandl, E. & Hojsgaard, D. 2012. The Evolution of Apomixis in Angiosperms: a reappraisal. Plant Biosystems, 146, 681–693.

Hu, T. T., Pattyn, P., Bakker, E. G., Cao, J., Cheng, J.-F., Clark, R. M., Fahlgren, N., Fawcett, J. A., Grimwood, J., Gundlach, H., Haberer, G., Hollister, J. D., Ossowski, S., Ottilar, R. P., Salamov, A. A., Schneeberger, K., Spannagl, M., Wang, X., Yang, L., Nasrallah, M. E., Bergelson, J., Carrington, J. C., Gaut, B. S., Schmutz, J., Mayer, K. F. X., Van De Peer, Y., Grigoriev, I. V., Nordborg, M., Weigel, D. & Guo, Y.-L. 2011. The Arabidopsis lyrata genome sequence and the basis of rapid genome size change. Nat Genet, 43, 476–481.

Huanca-Mamani, W., Garcia-Aguilar, M., Leon-Martinez, G., Grossniklaus, U. & Vielle-Calzada, J. P. 2005. CHR11, a chromatin-remodeling factor essential for nuclear proliferation during female gametogenesis in Arabidopsis thaliana. Proc Natl Acad Sci U S A, 102, 17231–6.

Huang, C. K., Huang, L. F., Huang, J. J., Wu, S. J., Yeh, C. H. & Lu, C. A. 2010. A DEAD-box protein, AtRH36, is essential for female gametophyte development and is involved in rRNA biogenesis in Arabidopsis. Plant Cell Physiol, 51, 694–706.

Hughes, S. C. & Simmonds, A. J. 2019. Drosophila mRNA Localization During Later Development: Past, Present, and Future. Front Genet, 10, 135–135.

Katoh, K., Misawa, K., Kuma, K.-I. & Miyata, T. 2002. MAFFT: a novel method for rapid multiple sequence alignment based on fast Fourier transform. Nucleic Acids Res, 30, 3059–3066.

Khanday, I., Skinner, D., Yang, B., Mercier, R. & Sundaresan, V. 2019. A male-expressed rice embryogenic trigger redirected for asexual propagation through seeds. Nature, 565, 91–95.

Kiefer, C., Willing, E. M., Jiao, W. B., Sun, H., Piednoël, M., Hümann, U., Hartwig, B., Koch, M. A. & Schneeberger, K. 2019. Interspecies association mapping links reduced CG to TG substitution rates to the loss of gene-body methylation. Nat Plants, 5, 846–855.

Kiefer, M., Nauerth, B. H., Volkert, C., Ibberson, D., Loreth, A. & Schmidt, A. 2020. Gene function rather than reproductive mode drives the evolution of RNA helicases in sexual and apomictic Boechera. Genome Biol Evol. 12, 656–573.

Klock, H. E. & Lesley, S. A. 2009. The Polymerase Incomplete Primer Extension (PIPE) method applied to high-throughput cloning and site-directed mutagenesis. Methods Mol Biol, 498, 91–103.

Kozlov, A. M., Darriba, D., Flouri, T., Morel, B. & Stamatakis, A. 2019. RAxML-NG: a fast, scalable and user-friendly tool for maximum likelihood phylogenetic inference. Bioinformatics, 35, 4453–4455.

Kumar, S. 2017. Epigenetic Control of Apomixis: A New Perspective of an Old Enigma. Adv Plants Agric Res, 7, 1–8.

Larsson, A. 2014. AliView: a fast and lightweight alignment viewer and editor for large datasets. Bioinformatics, 30, 3276–3278.

Lee, C. R., Wang, B., Mojica, J. P., Mandáková, T., Prasad, K., Goicoechea, J. L., Perera, N., Hellsten, U., Hundley, H. N., Johnson, J., Grimwood, J., Barry, K., Fairclough, S., Jenkins, J. W., Yu, Y., Kudrna, D., Zhang, J., Talag, J., Golser, W., Ghattas, K., Schranz, M. E., Wing, R., Lysak, M. A., Schmutz, J., Rokhsar, D. S. & Mitchell-Olds, T. 2017. Young inversion with multiple linked QTLs under selection in a hybrid zone. Nat Ecol Evol, 1, 119.

Li, H. 2013. Aligning sequence reads, clone sequences and assembly contigs with BWA-MEM. arXiv: Genomics.

Li, H. & Durbin, R. 2009. Fast and accurate short read alignment with Burrows-Wheeler transform. Bioinformatics (Oxford, England), 25, 1754–1760.

Li, H., Handsaker, B., Wysoker, A., Fennell, T., Ruan, J., Homer, N., Marth, G., Abecasis, G. & Durbin, R. 2009. The Sequence Alignment/Map format and SAMtools. Bioinformatics, 25, 2078–9.

Liao, Y., Smyth, G. K. & Shi, W. 2014. featureCounts: an efficient general purpose program for assigning sequence reads to genomic features. Bioinformatics, 30, 923–30.

Linder, P. & Owttrim, G. W. 2009. Plant RNA helicases: linking aberrant and silencing RNA. Trends Plant Sci, 14, 344–52.

Liu, M., Shi, D.-Q., Yuan, L., Liu, J. & Yang, W.-C. 2010. SLOW WALKER3, Encoding a Putative DEAD-box RNA Helicase, is Essential for Female Gametogenesis in Arabidopsis. J Integr Plant Biol 52, 817–828.

Liu, Y. & Imai, R. 2018. Function of Plant DExD/H-Box RNA Helicases Associated with Ribosomal RNA Biogenesis. Front Plant Sci, 9, 125

Mao, Y., Gabel, A., Nakel, T., Viehöver, P., Baum, T., Tekleyohans, D. G., Vo, D., Grosse, I. & Groß-Hardt, R. 2020. Selective egg cell polyspermy bypasses the triploid block. eLife, 9, e52976.

Mau, M., Lovell, J. T., Corral, J. M., Kiefer, C., Koch, M. A., Aliyu, O. M. & Sharbel, T. F. 2015. Hybrid apomicts trapped in the ecological niches of their sexual ancestors. Proc Nat Acad Sci, 112, E2357–E2365.

Mckenna, A., Hanna, M., Banks, E., Sivachenko, A., Cibulskis, K., Kernytsky, A., Garimella, K., Altshuler, D., Gabriel, S., Daly, M. & Depristo, M. A. 2010. The Genome Analysis Toolkit: a MapReduce framework for analyzing next-generation DNA sequencing data. Genome Res, 20, 1297–303.

Meinke, D. W. 2020. Genome-wide identification of EMBRYO-DEFECTIVE (EMB) genes required for growth and development in Arabidopsis. New Phytol, 226, 306–325.

Mingam, A., Toffano-Nioche, C., Brunaud, V., Boudet, N., Kreis, M. & Lecharny, A. 2004. DEAD-box RNA helicases in Arabidopsis thaliana: Establishing a link between quantitative expression, gene structure and evolution of a family of genes. Plant Biotechnol J, 2, 401–15.

Pagnussat, G., Yu, H.-J., Ngo, Q., Rajani, S., Sevugan, M., Johnson, C., Capron, A., Xie, L.-F., Ye, D. & Sundaresan, V. 2005. Genetic and molecular identification of genes required for female gametophyte development and function in Arabidopsis. Development (Cambridge, England), 132, 603–14.

Patterson, M., Marschall, T., Pisanti, N., Van Iersel, L., Stougie, L., Klau, G. W. & Schönhuth, A. 2015. WhatsHap: Weighted Haplotype Assembly for Future-Generation Sequencing Reads. J Comput Biol, 22, 498–509.

Pogorelko, G., Fursova, O. & Klimov, E. 2008. IDentification and Analysis of the Arabidopsis Thaliana Atfas4 Gene Whose Overexpression Results in the Development of A Fasciated Stem. J Proteomics Bioinform, 1, 329–335.

Posada, D. 2008. jModelTest: Phylogenetic Model Averaging. Mol Biol Evol, 25, 1253–1256.

Quinlan, A. R. & Hall, I. M. 2010. BEDTools: a flexible suite of utilities for comparing genomic features. Bioinformatics, 26, 841–2.

Rawat, V., Abdelsamad, A., Pietzenuk, B., Seymour, D. K., Koenig, D., Weigel, D., Pecinka, A. & Schneeberger, K. 2015. Improving the Annotation of Arabidopsis lyrata Using RNA-Seq Data. PLOS ONE, 10, e0137391.

Robinson, M. D., Mccarthy, D. J. & Smyth, G. K. 2009. edgeR: a Bioconductor package for differential expression analysis of digital gene expression data. Bioinformatics, 26, 139–140.

Ross, P., Slovin, J. & Chen, C. 2010. A simplifed method for differential staining of aborted and non-aborted pollen grains. Int J Plant Bioly, 1, 2.

Rotman, N., Durbarry, A., Wardle, A., Yang, W. C., Chaboud, A., Faure, J. E., Berger, F. & Twell, D. 2005. A novel class of MYB factors controls sperm-cell formation in plants. Curr Biol, 15, 244–8.

Schmid, M. W. 2017. RNA-Seq Data Analysis Protocol: Combining In-House and Publicly Available Data. In: Schmidt, A. (ed.) Plant Germline Development: Methods and Protocols. New York, NY: Springer New York.

Schmid Mw, S. A., Herrmann A, Hedhly A, Guthörl D, Grob S, Schmid P, Klostermeier UC, Rosenstiel P, Grossniklaus U. 2015. Polarized distribution of mRNA in the syncytial female gametophyte of Arabidopsis thaliana precedes cellularization and cell specification. PhD Theses, Universtiy of Zürich, Switzerland, http://www.zora.uzh.ch/id/eprint/113415/1/thesis_bot.pdf; chapter 3.

Schmid, M. W., Schmidt, A. & Grossniklaus, U. 2015. The female gametophyte: an emerging model for cell type-specific systems biology in plant development. Front Plant Sci, 6, 907.

Schmid, M. W., Schmidt, A., Klostermeier, U. C., Barann, M., Rosenstiel, P. & Grossniklaus, U. 2012. A powerful method for transcriptional profiling of specific cell types in eukaryotes: laser-assisted microdissection and RNA sequencing. PLoS One, 7, e29685.

Schmidt, A. 2020. Controlling Apomixis: Shared Features and Distinct Characteristics of Gene Regulation. Genes (Basel), 11, 329.

Schmidt, A., Schmid, M. W. & Grossniklaus, U. 2015. Plant germline formation: common concepts and developmental flexibility in sexual and asexual reproduction. Development 142, 229–241.

Schmidt, A., Schmid, M. W., Klostermeier, U. C., Qi, W., Guthorl, D., Sailer, C., Waller, M., Rosenstiel, P. & Grossniklaus, U. 2014. Apomictic and sexual germline development differ with respect to cell cycle, transcriptional, hormonal and epigenetic regulation. PLoS Genet, 10, e1004476.

Schmidt, A., Wuest, S. E., Vijverberg, K., Baroux, C., Kleen, D. & Grossniklaus, U. 2011. Transcriptome analysis of the Arabidopsis megaspore mother cell uncovers the importance of RNA helicases for plant germline development. PLoS Biol, 9, e1001155.

Schranz, M. E., Dobes, C., Koch, M. A. & Mitchell-Olds, T. 2005. Sexual reproduction, hybridization, apomixis, and polyploidization in the genus Boechera (Brassicaceae). Am J Bot, 92, 1797–810.

Shimizu, K. K., Ito, T., Ishiguro, S. & Okada, K. 2008. MAA3 (MAGATAMA3) Helicase Gene is Required for Female Gametophyte Development and Pollen Tube Guidance in Arabidopsis thaliana. Plant Cell Physiol, 49, 1478–1483.

Singh, L. N. & Hannenhalli, S. 2008. Functional Diversification of Paralogous Transcription Factors via Divergence in DNA Binding Site Motif and in Expression. PLOS ONE, 3, e2345.

Sizani, B. L., Kalve, S., Markakis, M. N., Domagalska, M. A., Stelmaszewska, J., Abdelgawad, H., Zhao, X. A., De Veylder, L., De Vos, D., Broeckhove, J., Schnittger, A. & Beemster, G. T. S. 2019. Multiple mechanisms explain how reduced KRP expression increases leaf size of Arabidopsis thaliana. New Phytol, 221, 1345–1358.

Sprunck, S. & Groß-Hardt, R. 2011. Nuclear behavior, cell polarity, and cell specification in the female gametophyte. Sex Plant Reprod, 24, 123–136.

Sprunck, S., Rademacher, S., Vogler, F., Gheyselinck, J., Grossniklaus, U. & Dresselhaus, T. 2012. Egg cell-secreted EC1 triggers sperm cell activation during double fertilization. Science, 338, 1093–7.

Stamatakis, A. 2006. RAxML-VI-HPC: maximum likelihood-based phylogenetic analyses with thousands of taxa and mixed models. Bioinformatics, 22, 2688–2690.

Stamatakis, A. 2014. RAxML version 8: a tool for phylogenetic analysis and post-analysis of large phylogenies. Bioinformatics, 30, 1312–1313.

Stein, R. E., Nauerth, B. H., Binmöller, L., Zühl, L., Loreth, A., Reinert, M., Ibberson, D. & Schmidt, A. 2021. RH17 restricts reproductive fate and represses autonomous seed coat development in sexual Arabidopsis. Development, 148, dev198739.

Tarazona, S., Furió-tarí, P., Turrà, D., Pietro, A. D., Nueda, M. J., Ferrer, A. & Conesa, A. 2015. Data quality aware analysis of differential expression in RNA-seq with NOISeq R/Bioc package. Nuc Acids Res, 43, e140–e140.

Tekleyohans, D. G., Nakel, T. & Groß-Hardt, R. 2017. Patterning the Female Gametophyte of Flowering Plants. Plant Physiol, 173, 122–129.

Terceros, G. C., Resentini, F., Cucinotta, M., Manrique, S., Colombo, L. & Mendes, M. A. 2020. The Importance of Cytokinins during Reproductive Development in Arabidopsis and Beyond. Int J Mol Sci, 21.

Twell, D., Oh, S.-A. & Honys, D. 2006. Pollen Development, a Genetic and Transcriptomic View. In: Malhó, R. (ed.) The Pollen Tube: A Cellular and Molecular Perspective. Berlin, Heidelberg: Springer Berlin Heidelberg.

Underwood, C. J., Vijverberg, K., Rigola, D., Okamoto, S., Oplaat, C., Camp, R., Radoeva, T., Schauer, S. E., Fierens, J., Jansen, K., Mansveld, S., Busscher, M., Xiong, W., Datema, E., Nijbroek, K., Blom, E. J., Bicknell, R., Catanach, A., Erasmuson, S., Winefield, C., Van Tunen, A. J., Prins, M., Schranz, M. E. & Van Dijk, P. J. 2022. A PARTHENOGENESIS allele from apomictic dandelion can induce egg cell division without fertilization in lettuce. Nat Genet, 54, 84–93.

Van Treeck, B. & Parker, R. 2018. Emerging Roles for Intermolecular RNA-RNA Interactions in RNP Assemblies. Cell, 174, 791–802.

Verkest, A., Weinl, C., Inzé, D., De Veylder, L. & Schnittger, A. 2005. Switching the cell cycle. Kip-related proteins in plant cell cycle control. Plant Physiol, 139, 1099–106.

Vijverberg, K., Ozias-Akins, P. & Schranz, M. E. 2019. Identifying and Engineering Genes for Parthenogenesis in Plants. Front Plant Sci, 10, 128.

Walden, N., German, D. A., Wolf, E. M., Kiefer, M., Rigault, P., Huang, X.-C., Kiefer, C., Schmickl, R., Franzke, A., Neuffer, B., Mummenhoff, K. & Koch, M. A. 2020. Nested whole-genome duplications coincide with diversification and high morphological disparity in Brassicaceae. Nat Commun, 11, 3795.

Wang, Z. P., Xing, H. L., Dong, L., Zhang, H. Y., Han, C. Y., Wang, X. C. & Chen, Q. J. 2015. Egg cell-specific promoter-controlled CRISPR/Cas9 efficiently generates homozygous mutants for multiple target genes in Arabidopsis in a single generation. Genome Biol, 16, 144.

Warnes, G., Bolker, B., Bonebakker, L., Gentleman, R., Huber, W., Liaw, A., Lumley, T., Mächler, M., Magnusson, A. & Möller, S. 2005. gplots: Various R programming tools for plotting data.

Willing, E.-M., Rawat, V., Mandáková, T., Maumus, F., James, G. V., Nordström, K. J. V., Becker, C., Warthmann, N., Chica, C., Szarzynska, B., Zytnicki, M., Albani, M. C., Kiefer, C., Bergonzi, S., Castaings, L., Mateos, J. L., Berns, M. C., Bujdoso, N., Piofczyk, T., De Lorenzo, L., Barrero-Sicilia, C., Mateos, I., Piednoël, M., Hagmann, J., Chen-Min-Tao, R., Iglesias-Fernández, R., Schuster, S. C., Alonso-Blanco, C., Roudier, F., Carbonero, P., Paz-Ares, J., Davis, S. J., Pecinka, A., Quesneville, H., Colot, V., Lysak, M. A., Weigel, D., Coupland, G. & Schneeberger, K. 2015. Genome expansion of Arabis alpina linked with retrotransposition and reduced symmetric DNA methylation. Nature Plants, 1, 14023.

Wuest, S. E., Vijverberg, K., Schmidt, A., Weiss, M., Gheyselinck, J., Lohr, M., Wellmer, F., Rahnenfuhrer, J., Von Mering, C. & Grossniklaus, U. 2010. Arabidopsis female gametophyte gene expression map reveals similarities between plant and animal gametes. Curr Biol, 20, 506–12.

Xu, R., Zhang, S., Huang, J. & Zheng, C. 2013. Genome-Wide Comparative In Silico Analysis of the RNA Helicase Gene Family in Zea mays and Glycine max: A Comparison with Arabidopsis and Oryza sativa. PLOS ONE, 8, e78982.

Yang, R., Jarvis, D., Chen, H., Beilstein, M., Grimwood, J., Jenkins, J., Shu, S., Prochnik, S., Xin, M., Ma, C., Schmutz, J., Wing, R., Mitchell-Olds, T., Schumaker, K. & Wang, X. 2013. The Reference Genome of the Halophytic Plant Eutrema salsugineum. Front Plant Sci, 4, 46.

Zhao, X., Bramsiepe, J., Van Durme, M., Komaki, S., Prusicki, M. A., Maruyama, D., Forner, J., Medzihradszky, A., Wijnker, E., Harashima, H., Lu, Y., Schmidt, A., Guthorl, D., Logrono, R. S., Guan, Y., Pochon, G., Grossniklaus, U., Laux, T., Higashiyama, T., Lohmann, J. U., Nowack, M. K. & Schnittger, A. 2017. RETINOBLASTOMA RELATED1 mediates germline entry in Arabidopsis. Science, 356, 6336.

Zühl, L., Volkert, C., Ibberson, D. & Schmidt, A. 2019. Differential activity of F-box genes and E3 ligases distinguishes sexual versus apomictic germline specification in Boechera. J Exp Bot, 70, 5643–5657.

