## Supporting Information for "Differential expression analysis of sexual and apomictic *Boechera* uncovers *FAS4* as crucial for gametogenesis"

### Supporting Information and supporting methods

### Supporting information

*FAS4 orthologues are active in reproductive tissues*

Expression analysis suggests potential roles of *FAS4* during gametogenesis. To gain further insights into the protein localization of FAS4 orthologues, we generated *A. thaliana* lines expressing *A. thaliana* FAS4 or the *B. stricta* orthologue in translational fusion to the genes encoding for the blue fluorescent protein PmTurquoise or the yellow fluorescent protein mVenus, respectively. Constructs were based on the genomic regions including upstream regulatory sequences (hereafter referred to as $AtFAS4_{genomic}$-*PmTurquoise* and *BsFAS4orth* $_{genomic}$-*mVenus*). Investigation of the lines identified *At*FAS4 activity in developing and mature female gametophytes and both orthologues in nuclei of anther tissues during pollen development (Figure S8).

*CRISPR/CAS9 lines devoid of the construct and harbouring a frame shift mutation could not be recovered*

The investigations on the lines generated by CRISPR/CAS9 were based on analysis of plants from T2 generation that have not been outcrossed to wild type and thus are likely to harbour at least one copy of the CRISPR/CAS9 expression cassette. For the lines under investigation, we identified alleles with frame shift insertions and putatively less deleterious in frame deletions (Table S9).

Due to the activity in egg cells or zygotes, additional defects during early embryogenesis could not fully be excluded. Nevertheless, earlier stages of gametogenesis should not to be affected by this. After crossing to wild-type, several plants devoid of the CRIPSR/CAS9 construct were obtained. But even though the parental lines harboured alleles with in frame deletions and frame shift insertions, from a total of 294 F1-offspring analysed only a few plants carrying a mutant allele were recovered, which only harboured in frame deletions (Table S9).

*CRISPR/CAS9 lines*

Plasmids pHEE401E harbouring sequences for small guiding RNAs were first transformed into the *E. coli* strain 10-beta (New England Biolabs) and confirmed by Sanger Sequencing at Eurofins Genomics (Eberbach, Germany) before transformation into the GV3101 chemically competent *Agrobacteria*. *Arabidopsis* plants were transformed by floral dip as described previously (Clough and Bent, 1998). Seeds of transformed plants were selected on hygromcycin. For genotyping the primers P1 and P2 were used to amplify the CRISPR/Cas9 target region by PCR with Phire Taq (Thermo Fisher Scientific) before restriction digestion of the PCR product H*ind*III (New England Biolabs). T2 plants that were offspring of two different T1 mother plants were used for investigation. Plants carrying a mutant allele were subsequently crossed to wild-type before selection of hygromycin sensitive plants. Absence of the T-DNA was further confirmed by PCR with primers P3 and P4. Primers used for genotyping were used to amplify the target region before cloning into pJET1.2 blunt (Thermo Fisher Scientific). Sanger sequencing with the T7 primer was used to identify the mutations using 6-8 clones per line. For backcrosses (Table S9) offspring of the lines were used to pollinate wild-type. 294 offspring of these crosses were selected and genotyped, and sequences were determined as described above.

*Confirmation of insertion site of T-DNA in fas4-1 and linkage to bar gene*

To confirm the insertion site of the T-DNA, flanking regions were amplified with primers LP and RB, and primers RP and LB to determine the insertion site of the right and left border of the T-DNA respectively. After amplification with Phire Hot Start II DNA Polymerase (Thermo Fisher Scientific) PCR products were cloned into pJET1.2 blunt (Thermo Fisher Scientific). Sanger Sequencing of the clones at Eurofins Genomics (Eberbach, Germany) using T7 primer confirmed the insertion in the 4th intron between bases 1210 and 1223 of the genomic DNA counted from the A of the start codon. Linkage of the bar gene conferring resistance to phosphinothricin was confirmed by genotyping of a large number of resistant plants.

*Generation of fluorescent marker lines for FAS4 orthologues*

Polymerase Incomplete Primer Extension (PIPE) cloning (Klock and Lesley, 2009) was applied for cloning of constructs for fluorescent markers of FAS4 activity. For cloning into linearized vectors with the Tmp-pSTB205 backbone (empty vectors or vectors harbouring *FAS4* constructs), linearized vectors and amplicons for inserts were transformed into chemically competent *E. coli*, either strain Dh5α (New England Biolabs) or Stbl2 (Thermo Fisher). First, a linear entry vector was generated by PCR amplification of Tmp-pSTB205 with primers P5 and P6. All linear vectors were amplified using PrimeStar GXL polymerase (Takara). Amplifications of genomic regions and coding sequences of fluorescent proteins were done using either Phusion or Q5 (New England Biolabs). The coding region of *AtFAS4* including the 3' UTRs was amplified from genomic DNA with primers P7 and P8. The amplified constructs were cloned into linear Tmp-pSTB205. To insert the upstream regulatory region into this clone, it was linearized and amplified with primers P9 and P10. The putative promoter region was amplified with primers P11 and P12 and integrated into the Tmp-pSTB205 vector the harbouring the coding region of *AtFAS4* to generate Tmp-pSTB205-AtFAS4. After validation of this clone by Sanger Sequencing (Eurofins Genomics, Eberbach), the coding region of PmTurquoise was inserted as C-terminal translational fusion construct. For this purpose, the Tmp-pSTB205-AtFAS4 was linearized by amplification with primers P13 and P14. The coding sequence of PmTurqouise was amplified from pGGD-PmTurquoise, kindly provided by Rainer Waadt (Centre for Organismal Studies (COS), Heidelberg University, Germany), with primers P15 and P16 and cloned into the linearized vector Tmp-pSTB205-AtFAS4 to generate Tmp-pSTB205-AtFAS4-PmTurqouise. For cloning the genomic region of the orthologue of *B. stricta FAS4* including the putative promoter region and the 3' UTR, on genomic DNA primers P17 and P18 were used. The amplified construct was integrated into the linearized Tmp-pSTB205 to generate Tmp-pSTB205-BsFAS4ortho. Afterwards, the coding sequence of mVenus was integrated to result in a C-terminal translational fusion construct. The mVenus coding sequence was amplified from pGGD-mVenus (Stein et al., 2021), kindly provided by Rainer Waadt (COS, Heidelberg University) with primers P19 and P20. Tmp-pSTB205-BsFAS4ortho was linearized with primers P21 and P22 to generate pSTB205-BsFAS4ortho-mVenus. The sequences of all clones were validated by Sanger Sequencing (Eurofins Genomics, Eberbach, Germany). Restriction digestion using P*vu*I (New England Biolabs) was applied before using Gateway LR clonase (Thermo Fisher) for cloning of the constructs into pEarlyGate301 to generate pEarlyGate201-AtFAS4-PmTurquoise and pEarlyGate201-BsFAS4ortho-mVenus, the expression constructs of *AtFAS4*$_{\text{genomic}}$-*PmTurquoise* and *BsFAS4orth* $_{\text{genomic}}$-*mVenus,* respectively.

*Clearing and microscopy*

Prior to clearing, buds and siliques were fixed in an ice-cold mixture of 75% ethanol and 25% acetic acid before vacuum infiltration 2x for 15 min on ice followed by overnight incubation in the fixative on ice before replacement of the fixative with 70 % ethanol. For clearing, buds and siliques were dissected with injection needles and treated with chloral hydrate/glycerol/water (8:1:2; w/v/v). For differential interference contrast (DIC) and epifluorescence microscopy, a Zeiss Axio Imager M1 (Zeiss, Oberkochen, Germany) or an Axioplan Imager (Zeiss) connected to a Leica DCM 2900 camera (Leica, Wetzlar, Germany) were used to capture the pictures. For epifluorescence microscopy ovules were mounted in 80% (w/v) glycerol and for laser scanning confocal microscopy in 5% glycerol with 0.1% (v/v) of the cell wall dye Renaissance 2200 (SR2200) (Musielak et al., 2015). For laser scanning confocal microscopy a Leica TCS SP8 microscope equipped with a 63x oil immersion objective and Leica LAS X software was applied to observe ovules from lines harbouring the p*AKV::H2B-YFP* marker. Excitations/emissions were set to 405/410-519 nm for SR2200, and 514/519-659 nm for YFP. Tissues harbouring the expression constructs *AtFAS4*$_{\text{genomic}}$-*PmTurquoise* and *BsFAS4orth* $_{\text{genomic}}$-*mVenus* were acquired sequentially with a Zeiss LSM710 microscope with excitations/emissions set to 405/453-564 and 514/519-605, respectively. For Alexander Stain and DAPI staining images were taken with a Zeiss Axioskop HBO50 connected to a Axiocam MrC5 or Axiocam HRm, respectively. Pictures were cropped and processed in Adobe Photoshop CS2 Version 9.0 (Adobe Systems Inc., San Jose, CA, USA) or ImageJ (https://imagej.nih.gov/ij/).

*Isolation of ovules and RNA for RNA-Seq*

To isolate ovules harbouring mature gametophytes, flower buds were emasculated. Three days after emasculation, ovules were isolated by manual microdissection with injection needles and immediately snap-frozen in liquid nitrogen. Minor contaminations with tissues from the surrounding siliques could not fully be avoided. Per sample, about 30-50 ovules pooled from 3 pistils were collected. Ovules were first ground using a Retsch Schwingmühle MM200 (Retsch, Germany), before RNA isolation with Arcturus PicoPure RNA Isolation Kit (Thermo Fischer Scientific) following manufacturer instructions for „Use with CapSure Macro LCM Caps", except that the extraction buffer was applied to the ground tissues. For all samples, RNA was treated with DNaseI (QIAGEN) on columns before elution with 11-15 µl of elution buffer.

*RNA quality control and amplification for RNA-Seq*

For all samples described in this study, RNA quality was determined using RNA Pico Chip on an Agilent 2100 Bioanalyzer (Agilent Technologies, USA). For linear amplification of RNA isolated from LAM dissected samples SMARTseq v4 Ultra Low Input RNA Kit for Sequencing (Takara Bio USA) was used following manufacturer instructions. Amplified cDNA was purified with AMPure Sample Purification Beads (Beckman Coulter, Brea, USA) and eluted in nuclease free $H_2O$.

*Germination and growth on medium supplied with streptomycin*

To test for potential association of FAS4 to ribosomes, we tested germination and growth of *fas4-1/FAS4* seeds and seedlings on Murashige and Skoog medium without streptomycin or supplied with 30 µg $ml^{-1}$ streptomycin. 14 days after plating of seeds which have previously been stratified for at least 1 day at 4 °C approximate rosette diameters were measured for a minimum of 9 seedlings for each genotype and condition. To test for significance of differences a two-sided students t-test was applied in R.

**References**

CLOUGH, S. J. & BENT, A. F. 1998. Floral dip: a simplified method for Agrobacterium-mediated transformation of Arabidopsis thaliana. *Plant J,* 16**,** 735-43.

KLOCK, H. E. & LESLEY, S. A. 2009. The Polymerase Incomplete Primer Extension (PIPE) method applied to high-throughput cloning and site-directed mutagenesis. *Methods Mol Biol,* 498**,** 91-103.

MUSIELAK, T. J., SCHENKEL, L., KOLB, M., HENSCHEN, A. & BAYER, M. 2015. A simple and versatile cell wall staining protocol to study plant reproduction. *Plant Reprod,* 28**,** 161-9.

STEIN, R. E., NAUERTH, B. H., BINMÖLLER, L., ZÜHL, L., LORETH, A., REINERT, M., IBBERSON, D. & SCHMIDT, A. 2021. RH17 restricts reproductive fate and represses autonomous seed coat development in sexual Arabidopsis. *Development,* 148**,** dev198739.
