## Supplemental Figures S1-S11 for "Differential expression analysis of sexual and apomictic *Boechera* uncovers *FAS4* as crucial for gametogenesis"

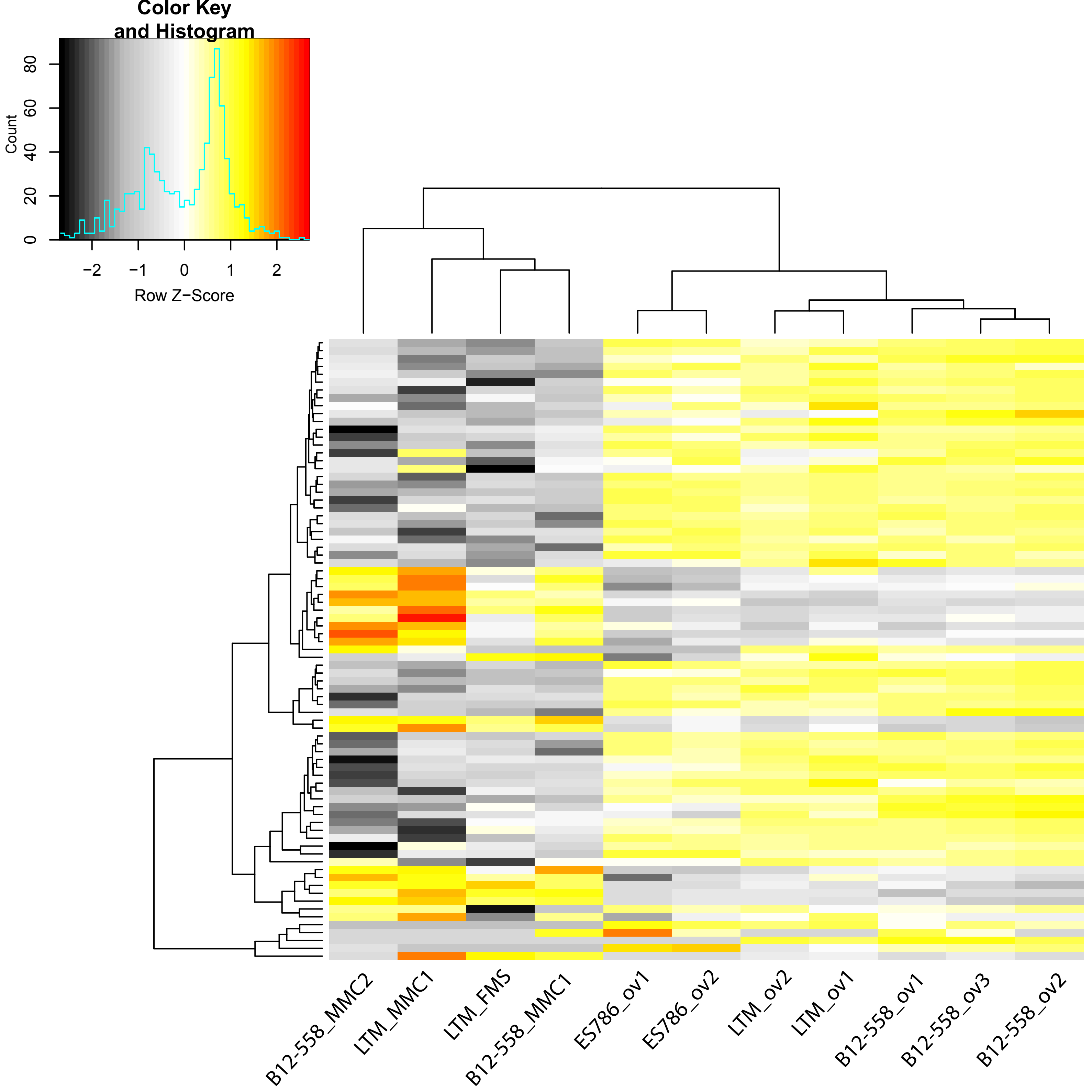


**Figure S1:** Heatmap of expression of 71 *Boechera* homologues of RNA helicases. The helicases were identified as DEGs in an ANOVA-like analysis comparing different stages of reproductive development in sexual accessions. The heatmap is based on log2-scale normalized read counts of all samples from sexual accessions of nucellus tissues harbouring the MMC, and of mature ovules. Hierarchical clustering of genes/samples was based on euclidian distance and hierarchical agglomerative clustering. Colors are scaled per row. Red denotes high and black low expression.

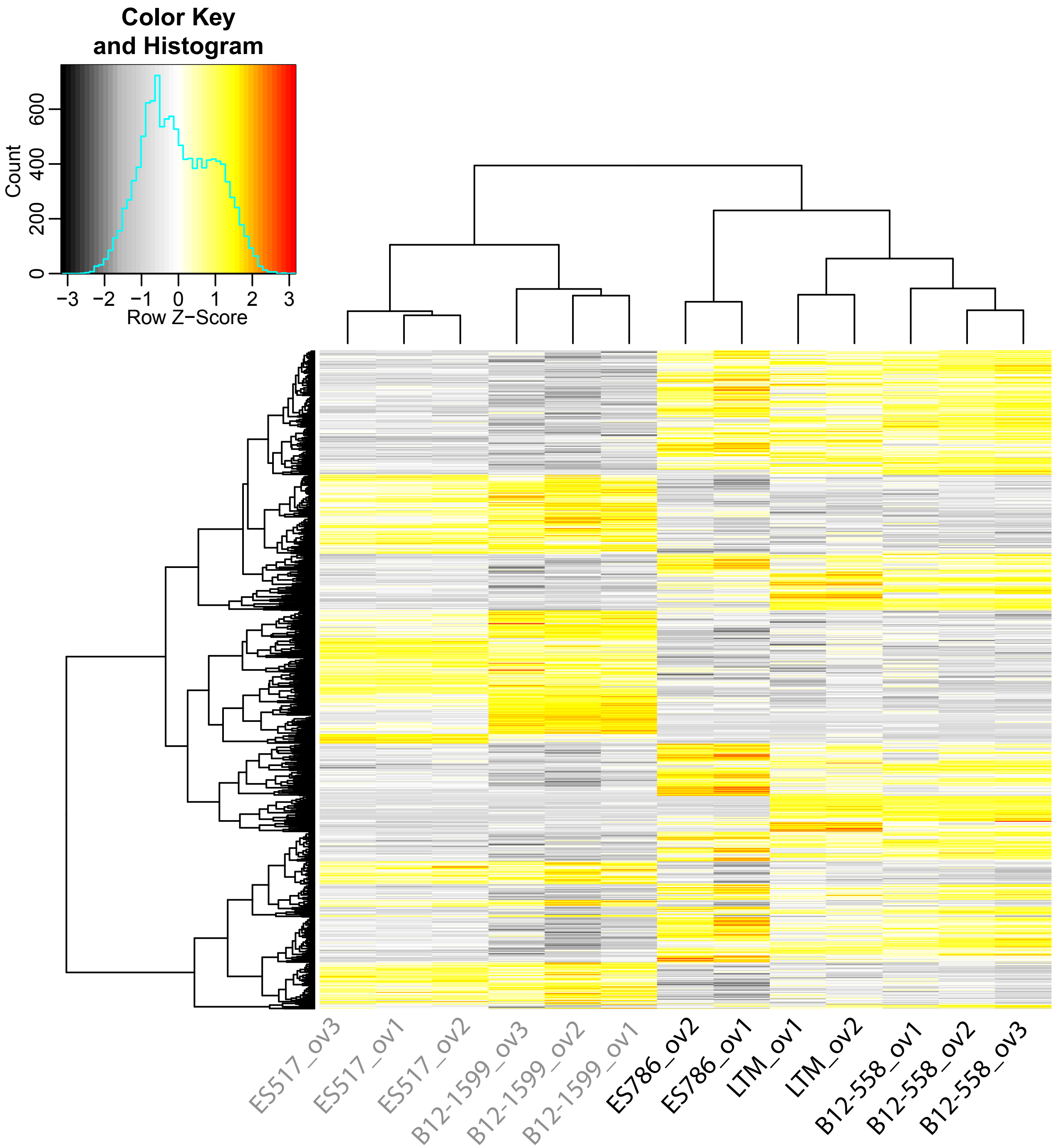


**Figure S2**: Heatmap of expression of 917 genes differentially expressed in ovules from sexual as compared to apomictic accessions. The heatmap is based on log2-scale normalized read counts. Hierarchical clustering of genes/samples was based on euclidian distance and hierarchical agglomerative clustering. Colors are scaled per row. Red denotes high and black low expression.

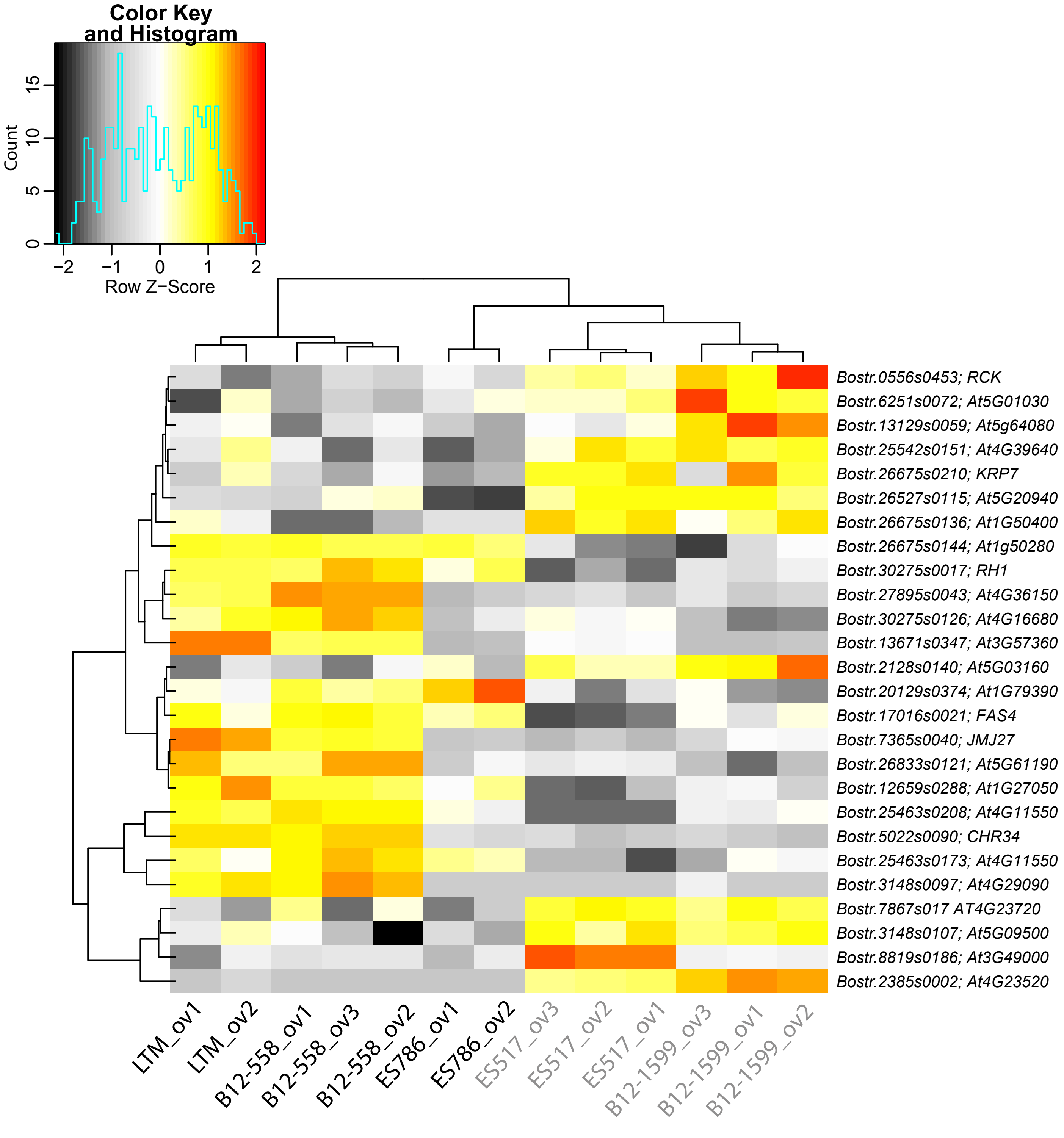


**Figure S3**: Expression of Arabidopsis gametophyte enriched genes and RNA helicases. Shown are such genes identified as DEGs in sexual as compared to apomictic ovules harbouring mature gametophytes. Gene identifiers are given for *B. stricta* LTM, in addition to gene names or identifiers of homologues in *A. thaliana*. The heatmap is based on log2-scale transformed TMM-normalized read counts. Hierarchical clustering of genes and samples was applied based on euclidean distance and hierarchical agglomerative clustering. Colors are scaled by row with red denoting high and black denoting low expression.

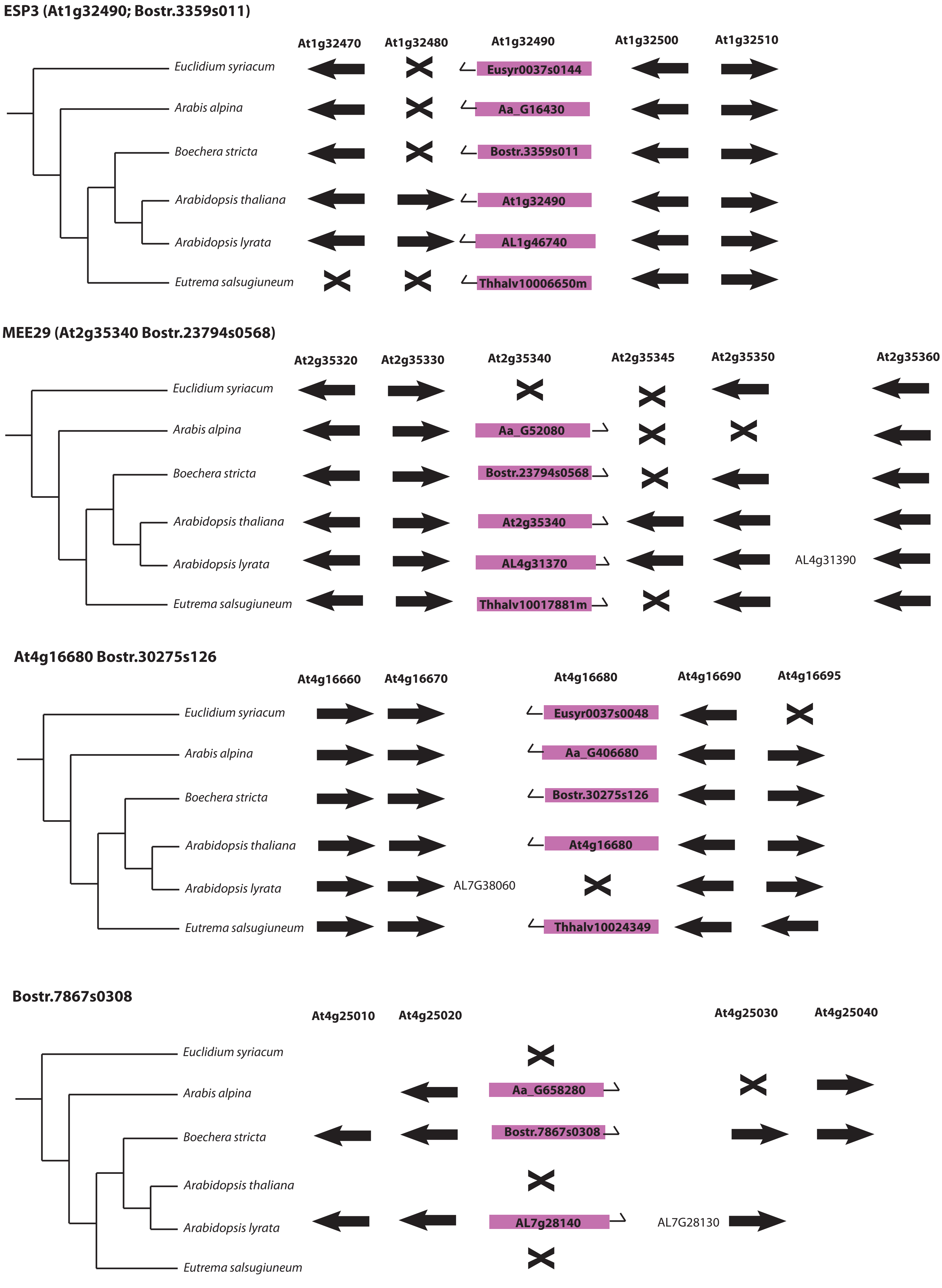


**Figure S4**: Evolutionary analysis of *ESP3*, *MEE29* and homologues identified *Bostr.3359s0111* as the positional orthologue of *AtESP3*, and *Bostr.30275s126* as orthologue of *At4G16680*, whereas no *A. thaliana* orthologue was found for *Bostr.7867s0308*.

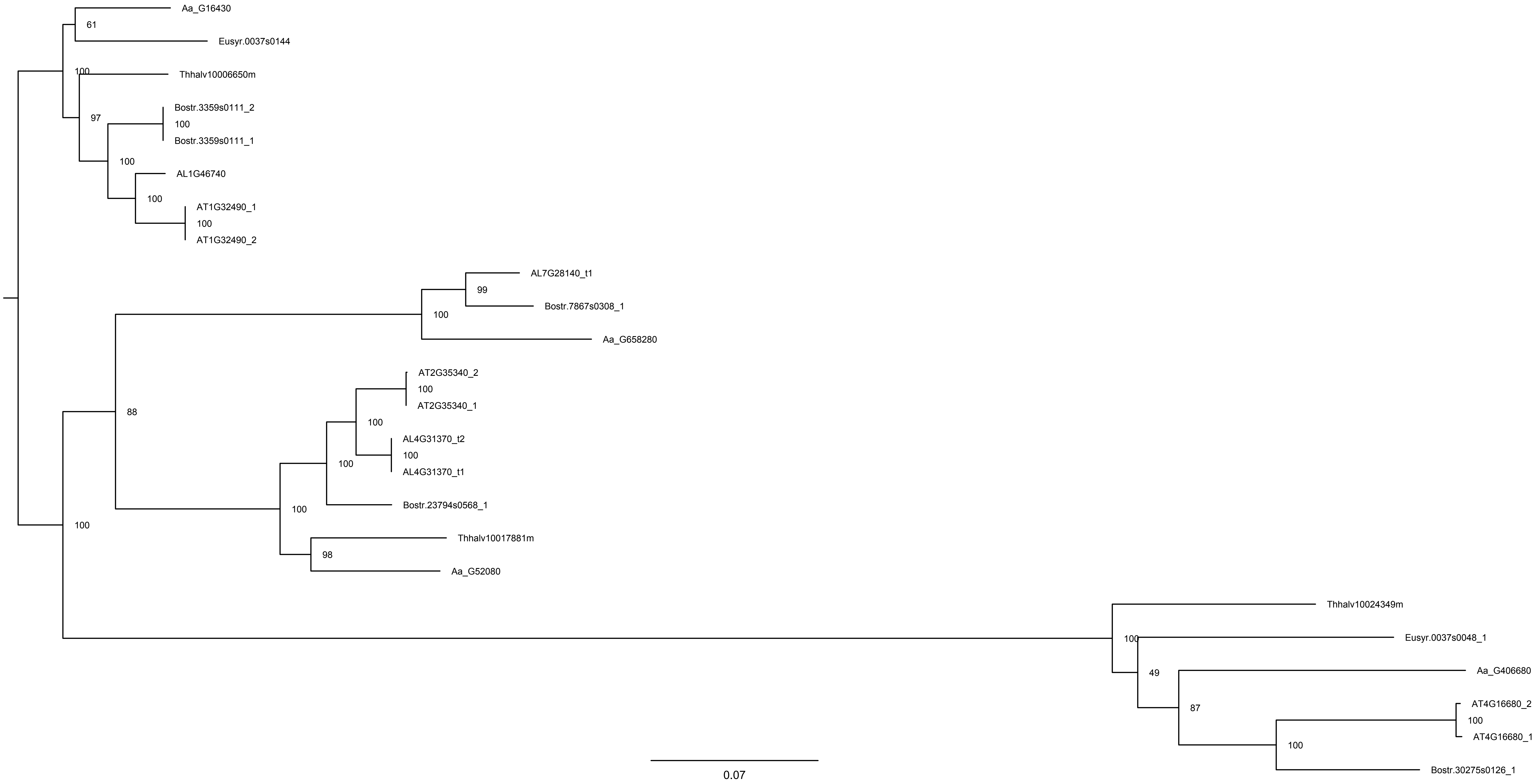


**Figure S5**: Phylogenetic analysis of homologues of *ESP3*, *MEE29* and related genes by maximum likelihood analysis using RAxML (Stamatakis, 2014).

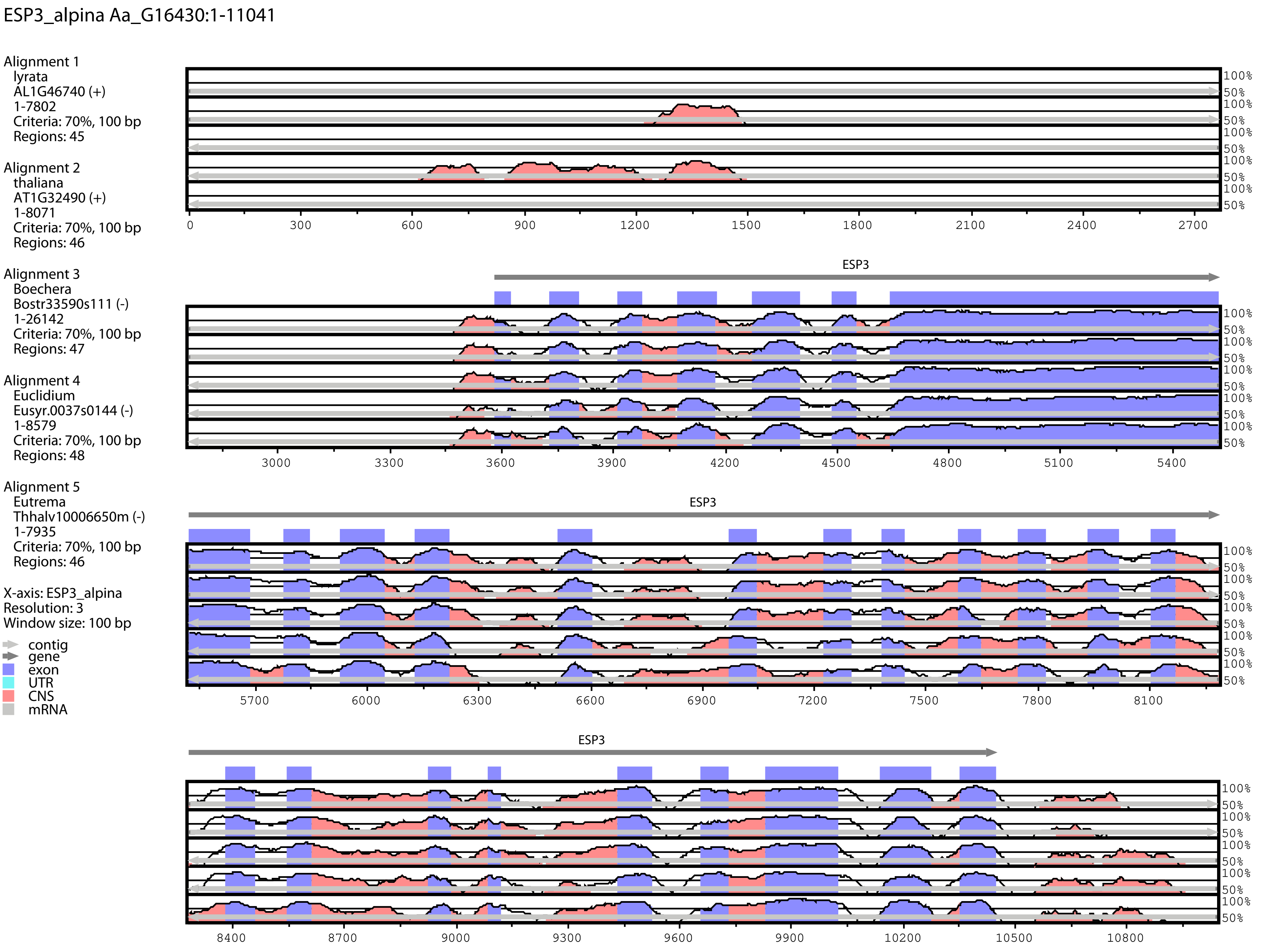
ESP3_alpina Aa_G16430:1-11041
Alignment 1
lyrata
AL1G46740 (+)
1-7802
Criteria: 70%, 100 bp
Regions: 45
Alignment 2
thaliana
AT1G32490 (+)
1-8071
Criteria: 70%, 100 bp
Regions: 46
Alignment 3
Boechera
Bostr33590s111 (-)
1-26142
Criteria: 70%, 100 bp
Regions: 47
Alignment 4
Euclidium
Eusyr.0037s0144 (-)
1-8579
Criteria: 70%, 100 bp
Regions: 48
Alignment 5
Eutrema
Thhalv10006650m (-)
1-7935
Criteria: 70%, 100 bp
Regions: 46
X-axis: ESP3_alpina
Resolution: 3
Window size: 100 bp
contig
gene
exon
UTR
CNS
mRNA
ESP3
100%
50%
0
300
600
900
1200
1500
1800
2100
2400
2700
3000
3300
3600
3900
4200
4500
4800
5100
5400
5700
6000
6300
6600
6900
7200
7500
7800
8100
8400
8700
9000
9300
9600
9900
10200
10500
10800

Figure S6A

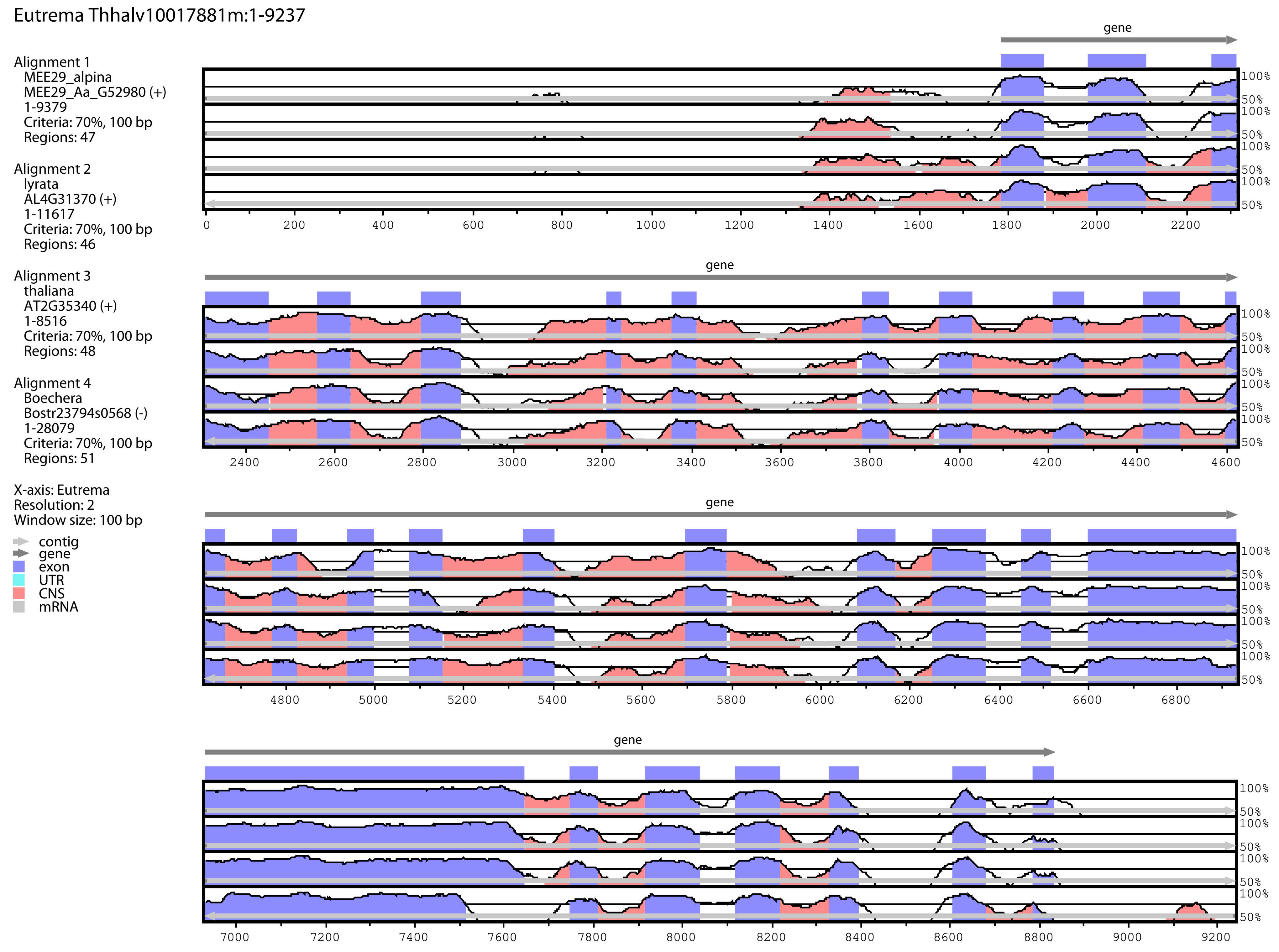
Eutrema Thhalv10017881m:1-9237
Alignment 1
MEE29_alpina
MEE29_Aa_G52980 (+)
1-9379
Criteria: 70%, 100 bp
Regions: 47
Alignment 2
lyrata
AL4G31370 (+)
1-11617
Criteria: 70%, 100 bp
Regions: 46
Alignment 3
thaliana
AT2G35340 (+)
1-8516
Criteria: 70%, 100 bp
Regions: 48
Alignment 4
Boechera
Bostr23794s0568 (-)
1-28079
Criteria: 70%, 100 bp
Regions: 51
X-axis: Eutrema
Resolution: 2
Window size: 100 bp
contig
gene
exon
UTR
CNS
mRNA
gene
100%
50%
0
200
400
600
800
1000
1200
1400
1600
1800
2000
2200
2400
2600
2800
3000
3200
3400
3600
3800
4000
4200
4400
4600
4800
5000
5200
5400
5600
5800
6000
6200
6400
6600
6800
7000
7200
7400
7600
7800
8000
8200
8400
8600
8800
9000
9200

Figure S6B

At4g16680_alpina Aa_G406800:1-8797

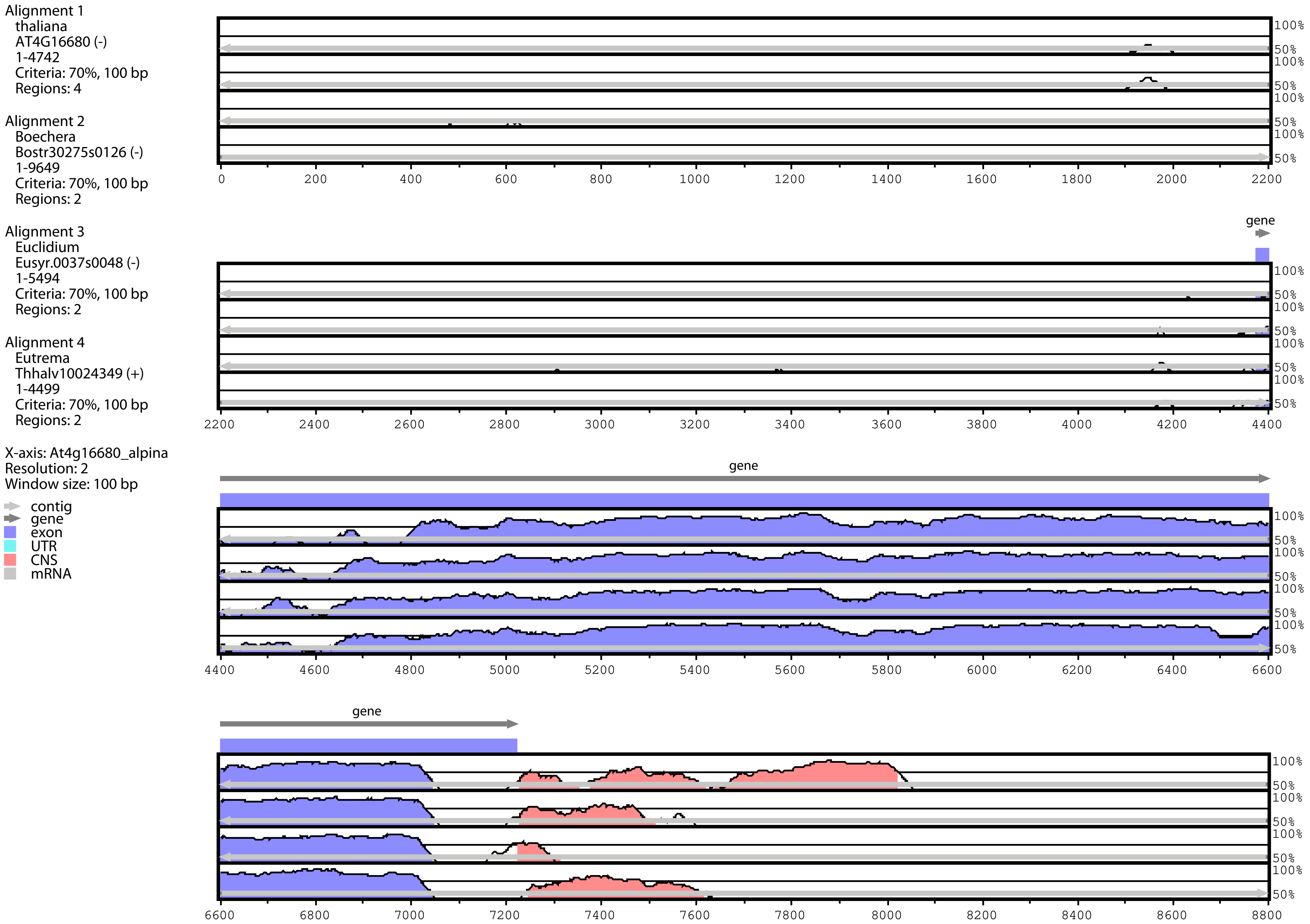


Figure S6C

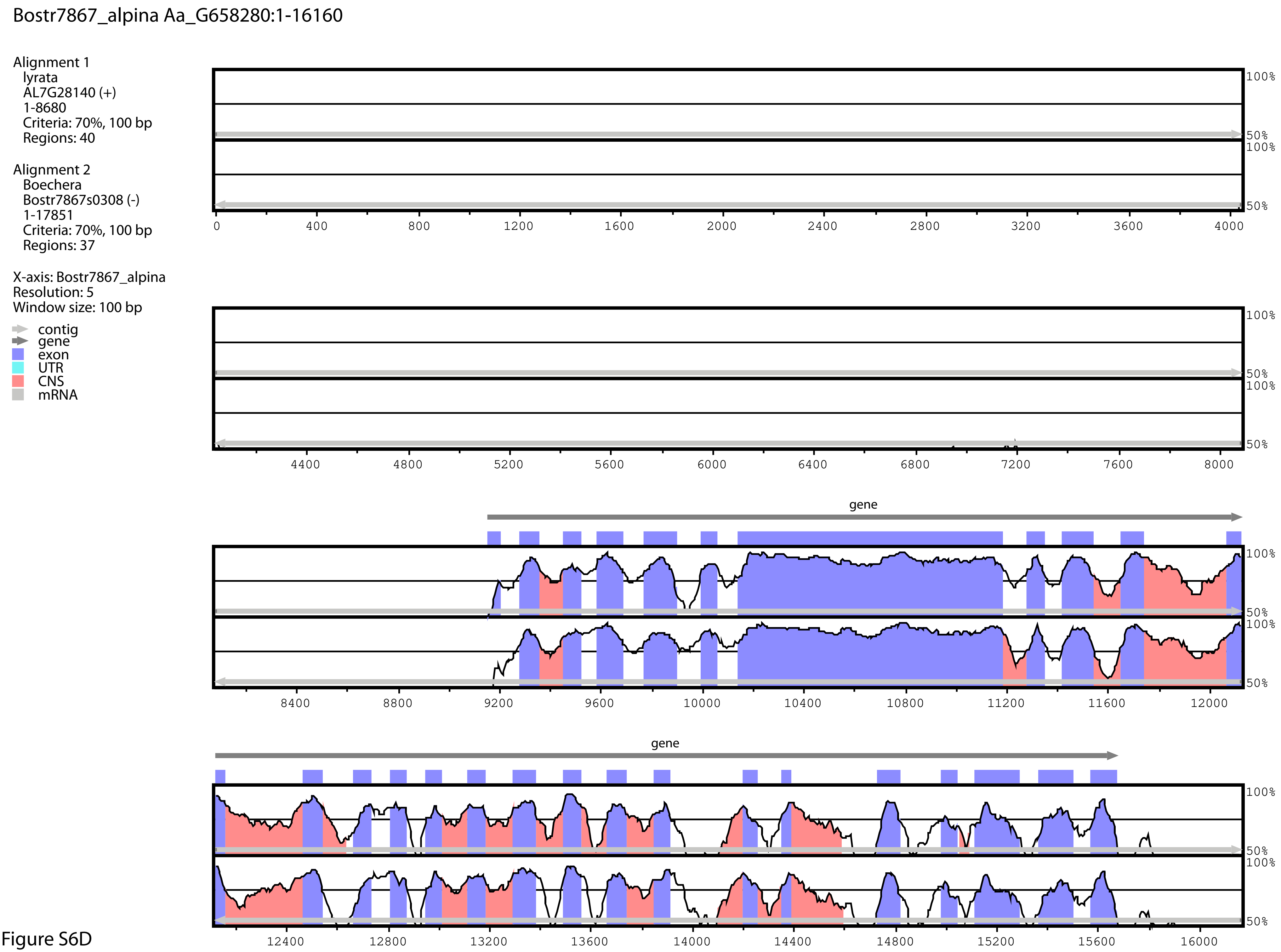


Figure S6D

**Figure S6**: Vista plots indicating similarities of orthologues from *ESP3* (A), *MEE29* (B), *At4G16680* (C), and *Bostr.7867s0308* in genomes of six selected Brassicaceae species using sequences from *A. alpina* (A,C,D) and *E. salsugineum* as base.

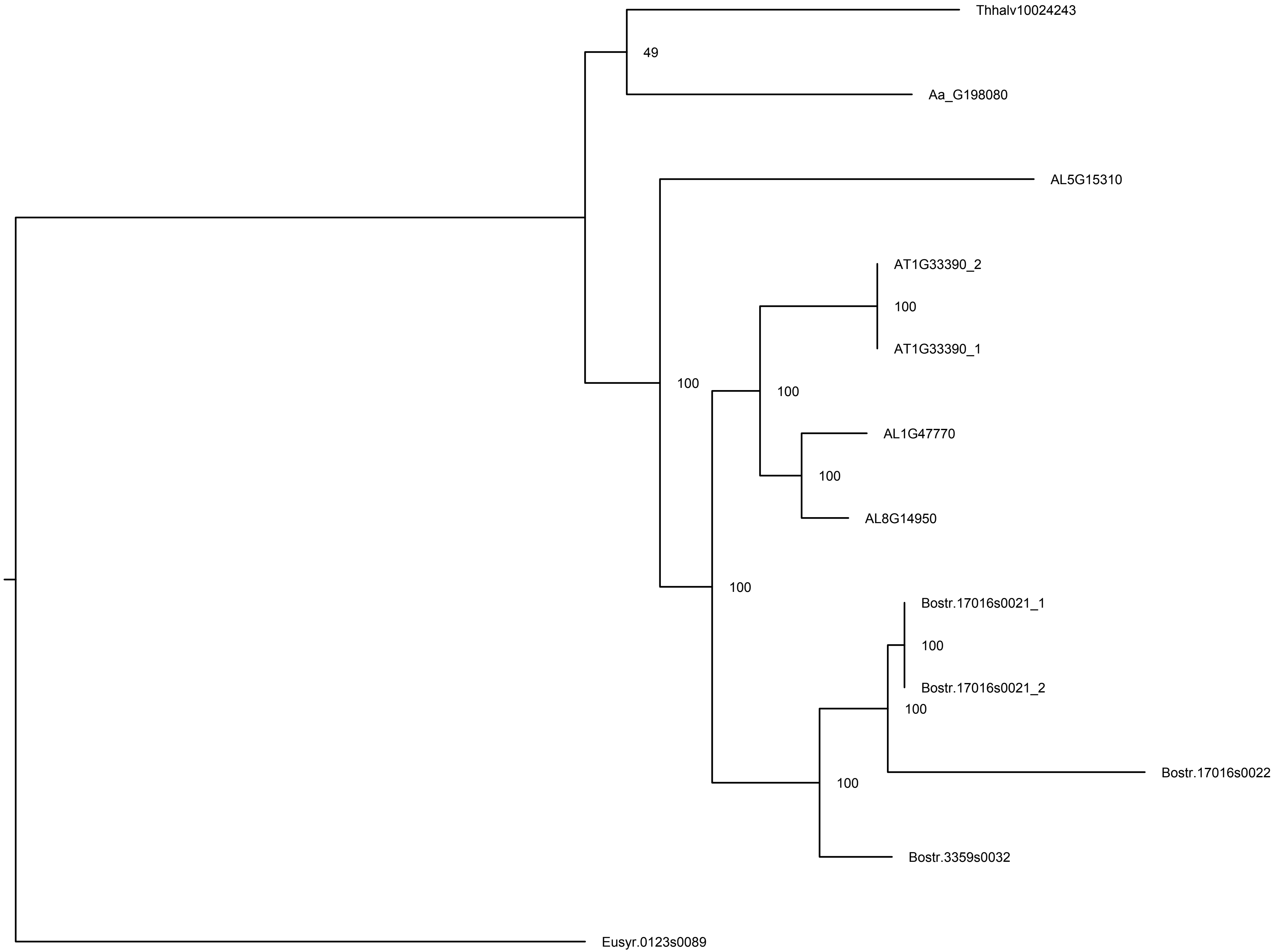


**Figure S7:** Phylogenetic analysis of homologues of *FAS4* by maximum likelihood analysis using RAxML (Stamatakis, 2014).

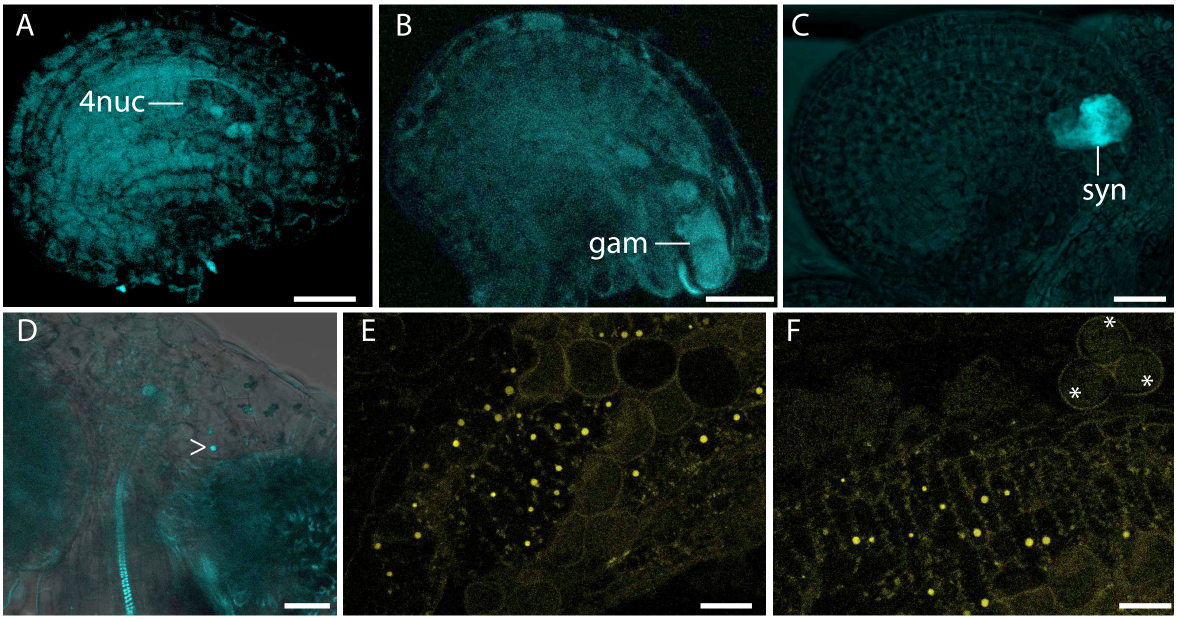


**Figure S8**: Localization of *At*FAS4 and *Bs*FAS4 orthologues in reproductive tissues using confocal laser scanning confocal microscopy. (A-D) Signals from fluorescence of PmTurquoise in plants carrying the *AtFAS4*genomic-*PmTurquoise* construct in developing ovule harbouring a developing gametophyte (A), mature gametophyte (B,C), and in anther tissues (D). (E,F) Signal from fluorescence of mVenus in anther tissues harbouring the and *BsFAS4orth*genomic-*mVenus*. 4nuc, 4 nucleate gametophyte; gam, mature gametophyte; syn, synergids; the arrow depicts a nucleus in anther tissues; * indicate pollen; scales = 20 µm.

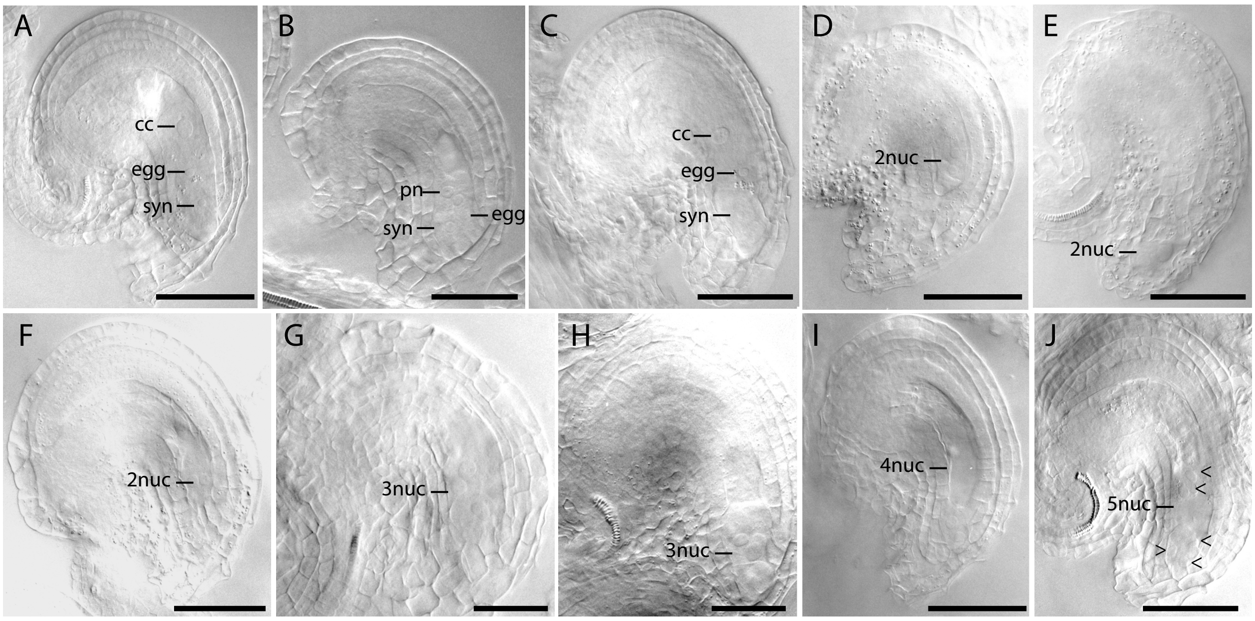


**Figure S9**: Clearing of *A. thaliana* ovules at 2 days after emasculation observed by DIC microscopy. (A,B) Wild-type ovules with mature gametophyte (A), and gametophyte with unfused polar nuclei (B). (C-J) Ovules from f*as4-1/FAS4* harbouring a wild-type mature gametophyte (C), an developmentally delayed or arrested 2-nucleate gametophyte (D), aberrant 2-nucleate gametophytes (E,F), a 4-nucleate gametophyte (I), and gametophytes with aberrant numbers of gametophytic nuclei (G,H,J). cc, central cell; egg, egg cell; syn, synergid cells; nuc, nucleate. Scale bars are 50 μm.

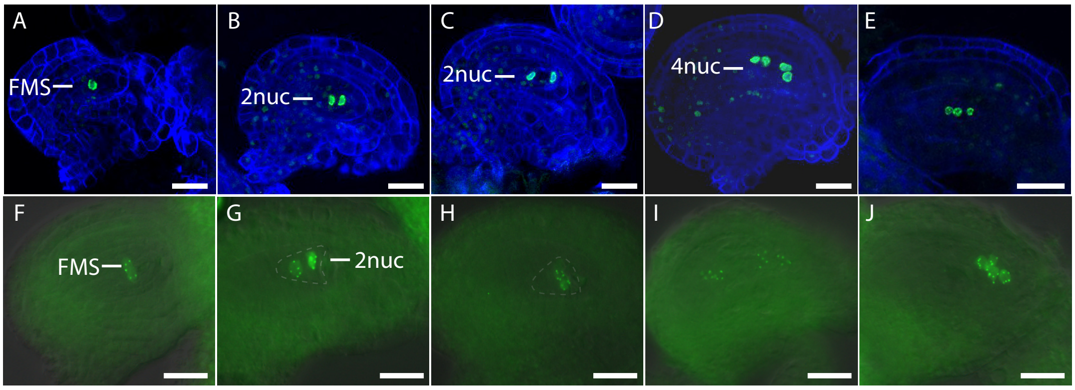


**Figure S10**: Developmental defects during female gametogenesis in *fas4-1/FAS4*. (A-E) Laser scanning confocal microscopy of developing ovules of plants harbouring the p*AKV::H2B-YFP* (Rotman et al., 2005) construct. Wild-type like FMS (A), 2-nucleate gametophyte (2nuc) at early (B) and later stage of development (C), and 4-nucleate gametophyte (4nuc) (D). (E) Mutant gametophyte showing an aberrant number of 3 gametophytic nuclei. (F-J) Epifluoresence microscopy on developing ovule carrying the p*WOX2::CENH3-GFP* marker (De Storme et al., 2016) for visualization of chromosomes in gametophytic nuclei. (F,G) Wild-type like gametophyte with 5 dots demarking the chromosomes in the haploid gametophytic lineage per nucleus in the FMS (F) and the 2-nucleate (2nuc) gametophyte (G). (H-J) Mutant gametophytes showing aberrations in the division plane/localization of gametophytic nuclei after the first mitotic division (H), alterations in migration (I), or separation of gametophytic nuclei (J). Scale bars = 20 μm.

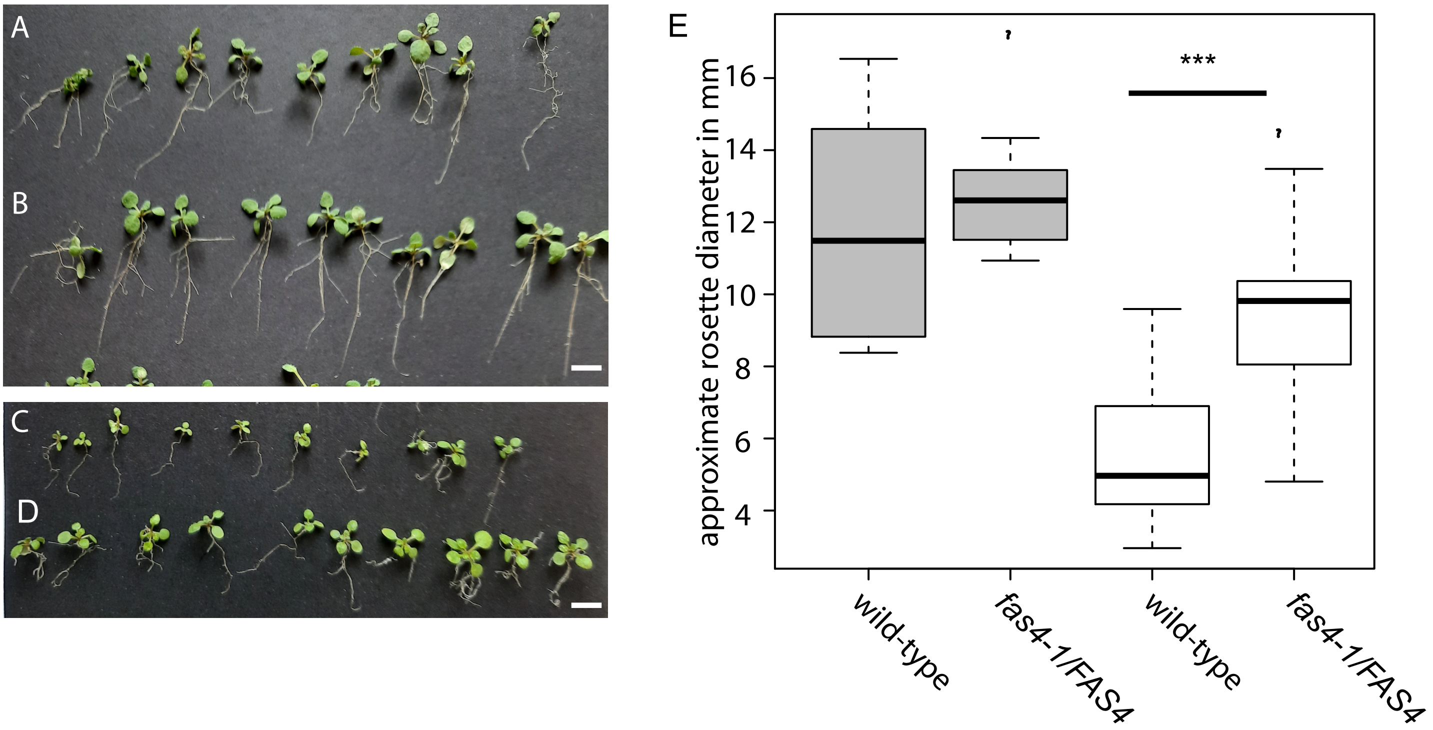


**Figure S11**: Seedlings of *fas4-1/FAS4* (B,D) show significantly increased resistance to streptomycin as compared to wild-type (A,C). (A-D) 14 day old seedlings germinated and grown on murashige-skoog medium without (A,B), or supplied with 30 µg ml-1 streptomycin (C,D). (E) Whereas no significant difference was observed concerning approximate rosette diameter in mm for 14 days old seedlings not supplied with streptomycin (gray boxes), on medium supplied with streptomycin rosette diameter were significantly larger for lines carrying a mutant allele of *FAS4* as compared to wild-type (white boxes, *** indicates $p < 0.005$ as tested with two-sided students t-test). Scales = 1 cm.
